# A single-cell atlas of bobtail squid visual and nervous system highlights molecular principles of convergent evolution

**DOI:** 10.1101/2022.05.26.490366

**Authors:** Daria Gavriouchkina, Yongkai Tan, Fabienne Ziadi-Künzli, Yuko Hasegawa, Laura Piovani, Lin Zhang, Chikatoshi Sugimoto, Nicholas Luscombe, Ferdinand Marlétaz, Daniel S. Rokhsar

## Abstract

Although the camera-type eyes of cephalopods and vertebrates are a canonical example of convergent morphological evolution, the cellular and molecular mechanisms underlying this convergence remain obscure. We used genomics and single cell transcriptomics to study these mechanisms in the visual system of the bobtail squid *Euprymna berryi*, an emerging cephalopod model. Analysis of 98,537 cellular transcriptomes from the squid visual and nervous system identified dozens of cell types that cannot be placed in simple correspondence with those of vertebrate or fly visual systems, as proposed by Ramón y Cajal and J.Z. Young. Instead, we find an unexpected diversity of neural types, dominated by dopamine, and previously uncharacterized glial cells. Surprisingly, we observe changes in cell populations and neurotransmitter usage during maturation and growth of the visual systems from hatchling to adult. Together these genomic and cellular findings shed new light on the parallel evolution of visual system complexity in cephalopods and vertebrates.

## Introduction

The camera eyes of vertebrates and cephalopods are a classical example of convergent evolution in two clades that diverged over 500 million years ago (dos Reis et al., 2015). Despite their optical parallels, however, cephalopod and vertebrate visual systems show marked differences in organization (Figure 1A). The vertebrate retina is a layered structure comprising multiple ciliary photoreceptors (rods and cones), oriented away from incoming light, whose output is processed through several layers of interneurons (amacrine, bipolar, and horizontal cells) before the visual signal is transmitted by ganglion cells to higher brain centers (Eakin, 1965; Shekhar and Sanes, 2021). In contrast, the cephalopod retina appears far simpler (Dilly et al., 1963) including only a single type of rhabdomeric photoreceptor oriented towards incoming light, and supporting cells but no interneurons. The cephalopod photoreceptor axons project directly from the retina to the large bilateral optic lobes of the central brain, which are organized into a layered outer cortex and inner medulla (Figure 1A,D-G).

**Figure 1.**
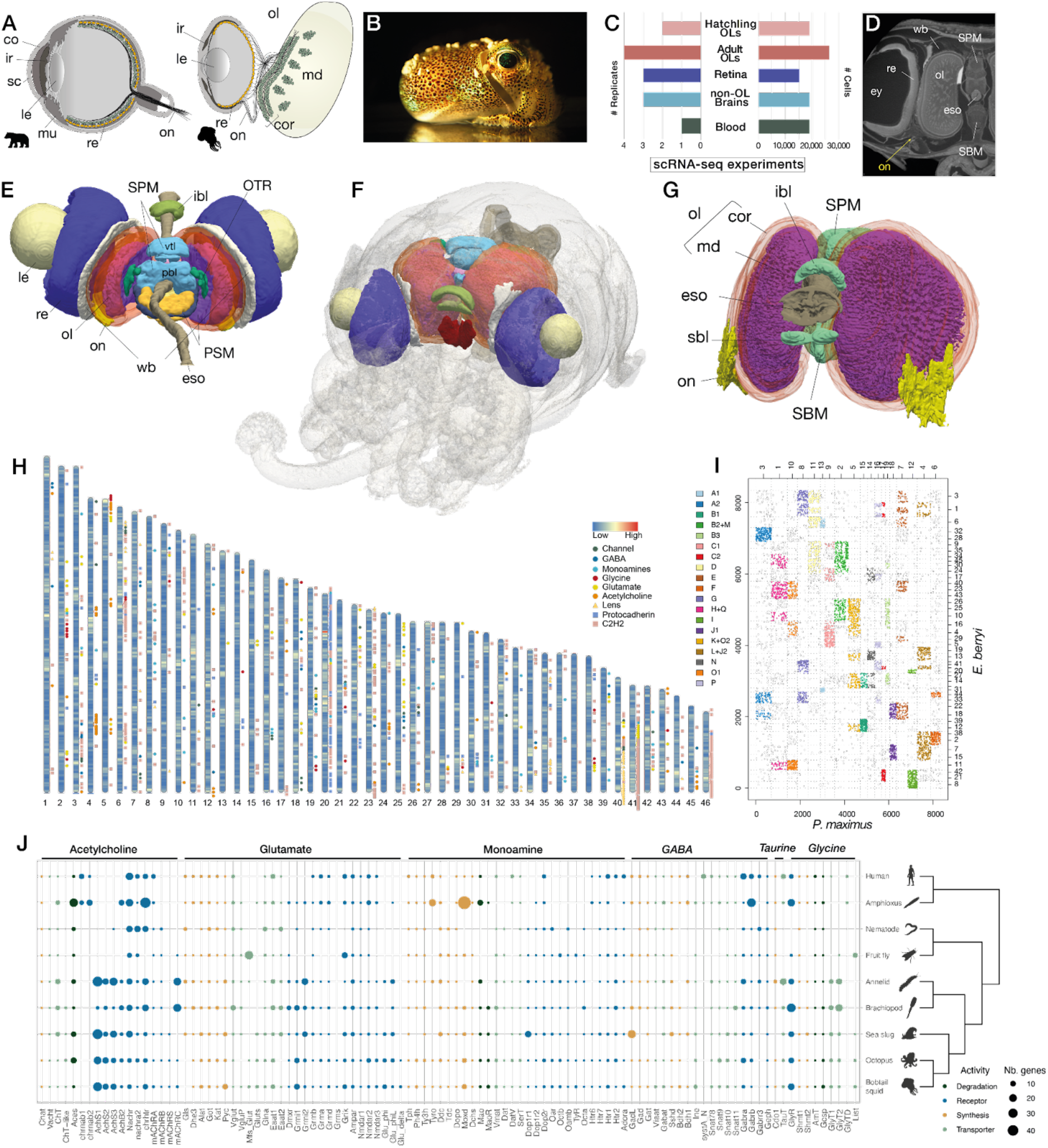
Genomic organization and neural gene complement of a cephalopod. (A) Simplified schematic representation of vertebrate and cephalopod camera-type eyes. Vertebrates (left) employ an inverted retina in which light traverses several layers of interneurons and ganglion cells (green) before reaching photoreceptor cells (yellow). Axons from the ganglion cells project to brain centers. The cephalopod retina (right) is believed to not feature any interneurons with light reaching the photoreceptors (yellow) directly. Axons from photoreceptors are thought to project onto the optic lobe cortex composed of cells that have been described as morphologically similar to those of the vertebrate interneurons in the retina by Santiago Ramón y Cajal and J.Z. Young. (B) Image of an adult *Euprymna berryi.* (C) Barplot indicating number of single cells sampled (right) and number of biological replicates (left). Number of technical replicates used is indicated in Supp. Data and Fig.S2A-D. (D-G) MicroCT characterization of the central nervous system of *E. berryi*. (D) Transverse section through microCT prior to rendering, indicating location of optic nerve in adults (70 dph) (yellow arrow). (E) MicroCT segmentation of the central nervous system of an adult *E. berryi* (70 dph) - postero-dorsal view, indicating eyes, brain lobes and associated white body. (F) MicroCT segmentation of the central nervous system of a hatchling *E. berryi, in toto* (1 dph) (G) Segmentation of adult optic lobe (ol) and substructures including cortex (cor) and medulla (md) - anterior view, oblique (70 dph). (H) Location of different categories of neural genes across the 46 chromosomes of *E. berryi*. (I) Oxford plot showing macrosyntenic blocks of orthologous genes located on *Euprymna berryi* chromosomes (y-axis) and a bivalve mollusc *Pecten maximus* (sea scallop) chromosomes (x-axis.) Positions along each axis represent integer-valued gene index. Color code (to the left) corresponds to ancestral linkage groups (ALGs).. Fusion events that took place in the cephalopod lineage can be observed at intersections of different ALGs. (J) Summary of phylogenetic analyses conducted for selected neural gene families and subfamilies stemming to the bilaterian ancestor involved in neurotransmitter synthesis, transport, degradation and reception. Monoamines include dopamine, serotonin, octopamine, and tyramine. Color code indicates activity (legend lower right). Species include human (*Homo sapiens*), amphioxus (*Branchiostoma lanceolatum*), nematode (*Caenorhabditis elegans*), fruit fly (*Drosophila melanogaster*), annelid worm (*Capitella teleta*), brachiopod (*Lingula anatina*), sea slug (*Aplysia californica*), octopus (*Octopus bimaculoides*) and bobtail squid (*Euprymna berry*i) with their relationships displays on the right. Abbreviations: ac - anterior chamber, co - cornea, cor - cortex, eso - esophagus, ey - eye, ibl - inferior buccal lobe, ir - iris, le - lens, md - medulla, mu - muscle, ol - optic lobe, on - optic nerve, PSM - posterior suboesophageal mass, re - retina, sbl - superior buccal lobe, SBM - subesophageal mass, ac - anterior chamber, SPM - supraesophageal mass, vbl - posterior basal lobe, vtl - vertical lobe, wb - white body.

Based on his pioneering observation of cell morphology and organization in the two systems, Santiago Ramón y Cajal proposed that the cephalopod optic lobe and vertebrate inner retina are functionally analogous (Ramón y Cajal, 1930; Young, 1962a, 1974). In particular, he recognized that in contrast to the tightly integrated system of the vertebrate retina, photoreception in the cephalopod retina is functionally separated from visual processing in the optic lobe. These considerations led him to suggest that the cortex of the cephalopod optic lobe, which he called ‘the deep retina’ (‘*la retina profunda*’) is analogous to the vertebrate neural retina. This picture gained further support from the detailed morphological analyses of J. Z. Young, who further characterized the morphological cell types and connectivity of the octopus and squid visual systems, including feedback from the optic lobe to the photoreceptor (Ramón y Cajal, 1930; Young, 1962a, 1974).

Evaluating the ‘deep retina’ hypothesis raises fundamental questions about the evolution/homology of complex systems. In its most literal sense, this hypothesis suggests a deep evolutionary correspondence between cell types of the vertebrate retina and the cephalopod optic lobe. We now understand, based on information unavailable to Ramón y Cajal, that these two complex visual systems evolved convergently (Ogura et al., 2004). Direct cell type homology (in the sense of cell types in modern cephalopod and vertebrate eyes descending from specific ancestral cell types in their common ancestor (Arendt et al., 2016)) is therefore unlikely, since the last common ancestor of molluscs and chordates did not have an image forming eye (Nilsson, 2009; Salvini-Plawen et al., 1977). Instead, we must consider that convergent structures could arise by some combination of (1) parallel reuse of shared ancestral genes or networks (Shubin et al., 2009), perhaps following common organizational principles (Sanes and Zipursky, 2010) and (2) the parallel acquisition of new genes and pathways, such as novel adhesion molecules or transcriptional activators (Albertin et al., 2015a, 2022b). From this perspective, the *a priori* convergent cephalopod and vertebrate visual systems provide an ideal opportunity to begin to compare the genes and cell types underlying convergently evolved complex systems. Although more than 60% of transcripts expressed in the octopus eye have orthologs that are also expressed in vertebrate eyes (Ogura et al., 2004), we do not know how these genes are deployed at the cellular level, or the role of gene novelty in structural and functional convergence.

To this end, we use single-cell transcriptomics (scRNA-seq) to characterize the visual system and central nervous system of the Japanese bobtail squid *Euprymna berryi* Sasaki 1929 (Sasaki, 1929), and compare the cellular composition and organization of squid and vertebrate visual systems. *E. berryi* is an emerging cephalopod model that is closely related to other bobtail squids (McFall-Ngai, 2008; Nabhitabhata and Nishiguchi, 2014), but is somewhat larger and more amenable to laboratory culture. MicroCT reconstruction confirms that the *E. berryi* visual system bears the classic cephalopod organization (Chung et al., 2020; Liu et al., 2017; Wild et al., 2015) (Figure 1D-G). For the purposes of analyzing single cell data, a reference genome better captures the complexity of cephalopod gene expression than *de novo* assembled transcriptomes, so we generated a high-quality chromosome-scale genome assembly and annotation for *E. berryi* (Figure 1H-J).

Our collected datasets and analyses shed new light on the cephalopod visual system and represent first steps toward a molecular and cellular understanding of cephalopod visual systems. We confirm the simplicity of the squid retina, but find two photoreceptor sub-types distinguished by the expression of two rhabdomeric opsins (r-opsin paralogs). In contrast we observe a far greater diversity of cell types in the optic lobe, with diverse neurotransmitters dominated by dopamine. Both optic lobes and retina employ an extensive repertoire of neuropeptides, and we find that FMRF serves as the retrograde signal hypothesized by Young (Young, 1962a). We identify putative glial cells, filling a gap in our understanding of cephalopod nervous systems (Ibrahim et al., 2020; Imperadore et al., 2017). Surprisingly, the cell complements of one-day old hatchlings and mature adults differ, with considerably fewer dopaminergic cells in the hatchlings, consistent with changes in optic lobes seen with microCT during maturation and observations in cuttlefish (Liu et al., 2017). Cross-species comparisons of cell type expression between bobtail squid, vertebrates, and flies show that cell types cannot be placed in simple correspondence except for coarse relationships based on neurotransmitter usage. We find that genes involved in neuronal cell types have a complex evolutionary history, including neurotransmitter receptors, cephalopod-specific transcription factors and adhesion molecules, which participated in the independent origins of neural cell types in diverse animals.

## Results

### A genomic resource for an amenable bobtail squid model

As a foundation for single-cell transcriptomics, we assembled a highly contiguous chromosome-scale reference genome for the Japanese bobtail squid *E. berryi* by combining long-read sequencing with chromatin conformation capture data (Figure 1H, Figure S1A-B, Table S1). Our assembly of the *E. berryi* genome totals 5.5 Gb and captures the 46 chromosomes found in decapods in a more contiguous fashion (N50 chromosome length: 113.96 Mb, N50 contig length: 827 kb) than other available cephalopod genomes (Albertin et al., 2022a, 2015b; Belcaid et al., 2019; Kim et al., 2018; Zarrella et al., 2019; Zhang et al., 2021). For instance, we recovered the massive 17 Mb Hox cluster as an intact locus on chromosome 9 (Figure S1G). Different expansions of LINE transposons and other repeats may explain the difference in genome size as compared to other cephalopod species (Figure S1C). To support single cell and comparative analysis, we generated a combination of bulk short-read and long-read RNA-seq (Iso-seq) from diverse neural and non-neural tissues, as well as a range of whole animal embryonic stages (Table S2). We used this transcriptome data to annotate 32,244 protein-coding genes in *E. berryi,* with detectable homology in other animals.

*E. berryi* chromosomes are generally derived from the fusion of several ancestral linkage groups (ALGs) as shown by the comparison with the bivalve mollusc *Pecten maximus*: for instance, the tandem array-rich chromosome 20, corresponds to a mixture of ALGs I and G (Albertin et al., 2022b; Simakov et al., 2022) (Figure 1H-J). The karyotype of cephalopods is heavily affected by such fusion events and conceals multiple tandem expansion hotspots, whose respective contribution to the emergence of their unique features appears essential (Figure 1H-J). However, the 46 chromosomes of *E. berryi* shows extensive conservation with the distantly related *D. pealei* the longfin inshore squid (Figure S1D) (Albertin et al., 2022c).

The *E. berryi* genome encodes an extensive complement of genes associated with neurotransmitter synthesis, degradation, transport and reception (Figure 1H,J). We assessed evolutionary trends in these neuronal genes across bilaterians using gene family reconstruction and phylogenetic analysis (Methods). While genes associated with neurotransmitter biosynthesis and degradation have remained in relatively stable copy numbers across bilaterians, we find extensive lineage-specific expansions in neurotransmitter receptors and transporters in *E. berryi* and other spiralians (Figure 1J). In general, we see more variation in copy number of genes associated with acetylcholine and glutamate activity, whereas inhibitory GABA and glycine activity appears more uniform in copy number across bilaterians. For example, we identified single copies of monoaminergic synthesizing enzymes such as tyrosine hydroxylase (Ty3h), tryptophan hydroxylase (Tph2) or tyrosine decarboxylase (Tdc1) across bilaterians. In contrast, the *E. berryi* genome encodes multiple ionotropic acetylcholine receptor subunits, as well as multiple spiralian-specific families of glutamate receptors, both metabotropic (e.g., Grma-Grmi) and ionotropic (e.g., GluPhi, Phi-like and Delta) (Jiao et al., 2019; Ramos-Vicente et al., 2018). The absence of the glutaminase enzyme in *E. berryi* and other cephalopods suggests that glutamate synthesis is accomplished through another mechanism, possibly using Krebs cycle derivatives.

Expanded gene families found in *E. berryi* are typically found in gene duplication hotspots, which are extensive across the genome (Figure 1H). For example, *E. berryi* shares an expansion of ionotropic acetylcholine receptors described in lophotrochozoans (Jiao et al., 2019) located in two main hotspots on chromosomes 4 and 5. Interestingly, parallel expansions occurred independently in other spiralian lineages, such as the annelid *Capitella teleta* and the gastropod mollusc *Aplysia californica* (Figure 1J, Figure S1). We found extensive tandem arrays of C2H2 transcription factors on chromosomes 3, 6, 41, 46 and an expansion of both C2H2 transcription factors and protocadherins, on chromosome 20, paralleling findings in other cephalopod species (Albertin et al., 2022b) (Figure 1J). Moreover, the cephalopod lens is composed of glutathione-S-transferase proteins known as σ-crystallins expanded in 193 copies in two hotspots on chromosomes 40 and 41.

*E. berryi* preserves the ancestral bilaterian complement of neural genes better than other lineages including, for example, sub-families that have been lost in ecdysozoans or vertebrates (Figure 1J). Some of these sub-families are only present in spiralians, including spiralian-specific ionotropic acetylcholine receptor subunit families (AchS1-AchS3), metabotropic acetylcholine receptors (mAchRC), metabotropic glutamate receptor families (Grmi1-2 and Grms), octopamine and tyramine receptors (Oar, Octa, Octb, Oamb, TyR families), serotonin receptors (Htr6 family) or inhibitory neurotransmitter transporter GlyT2 (Figure 1J). However, it is noteworthy that other spiralians, such as annelids and lophophorates, have similarly diverse neural gene complements, which suggests that gene diversification alone is insufficient to explain cephalopod neural complexity.

### A single cell atlas of the squid visual system

To investigate possible cellular correspondences between cephalopods and vertebrates, we dissected components of the squid visual system and applied droplet-based scRNA-seq to characterize cell type-specific gene expression (Figure 1C). In total, we characterized the transcriptomes of 98,537 cells, including the adult retina (15,223 cells), optic lobes (19,029 cells from one day old hatchlings and 26,436 cells from mature adults) and non-visual organs (17,468 cells from non-optic lobe adult brain cells and 20,381 blood cells from mature adults) (Figure 1C, Figure S2A-D). Neurons were identified as cells positive for neuronal markers synaptotagmin (Sy65), voltage-dependent calcium channel subunit alpha-1 (Cac1a), and a voltage-gated sodium channel (Scna) (Figure 2B, Figure 3B). Putative glial cells were identified as described below. To begin to determine the spatial distribution of cell-types in the visual system, we used hybridization chain reaction (HCR) to locate mRNAs of selected cell-type-specific markers on both whole-mount preparations and sections of individual *E. berryi* (10 dph) (Choi et al., 2018) (Figure 2D-G, Figure 3E-L’, Figure S4E-F, Figure 5G-J).

**Figure 2.**
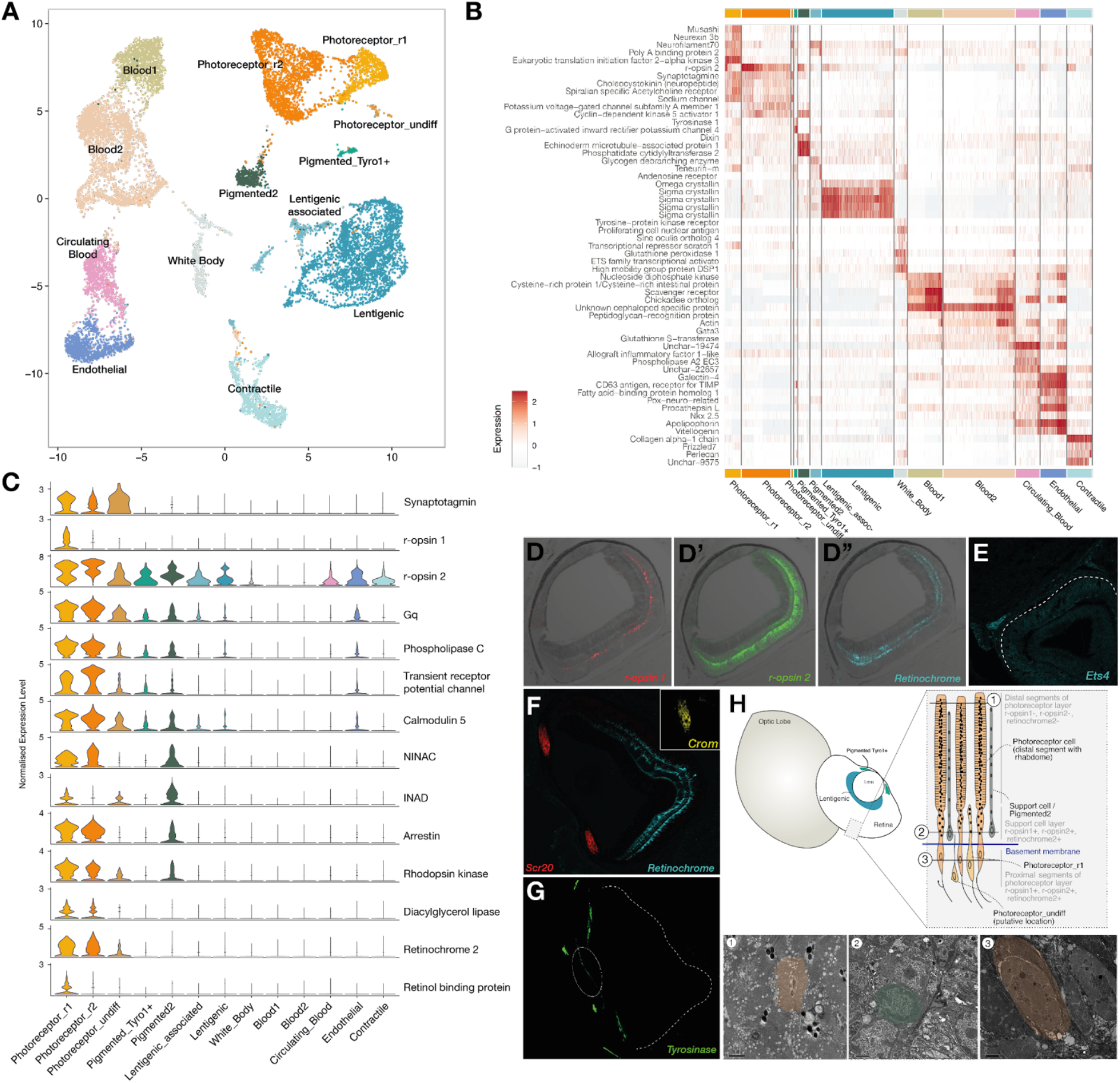
Cell type complement of the *E. berryi* retina. (A) UMAP of 15,223 cells from dissected adult *E. berryi* retinas (B) Heatmap indicating genes specifically expressed in each set of cell clusters. (C) Violin plots indicating expression levels of genes involved in phototransduction of rhabdomeric opsins and visual cycle. (D-G) Fluorescence imaging of HCR stainings: (D-D’’) three photoreceptive molecules r-opsin1, r-opsin2 and retinochrome2, displaying expression demonstrating localisation of r-opsin1 in ‘support cell layer’ and r-opsin2 and retinochrome-2 in support cell layer and in the proximal segment layer of the retina. (E) Ets4 expression in the presumptive white body between the retina and optic lobe (Yoshida et al., 2015) (F) retinochrome-2 and two lentigenic markers σ-crystallin (Scr20-109) and Ω-crystallin (Crom). (G) Tyrosinase (Tyro) expression in tissues surrounding the pupil of the retina. (H) Schematic of retina indicating putative location of cell clusters within retinal organization and images of a section through the retina at the level of the support cell layer, demonstrating the presence of multiple cell types at levels indicated in the schematic demonstrating 1- rhabdomes of photoreceptor cells, 2- Support cells near basement membrane, 3- Proximal segments of photoreceptor cells featuring nuclei.

**Figure 3.**
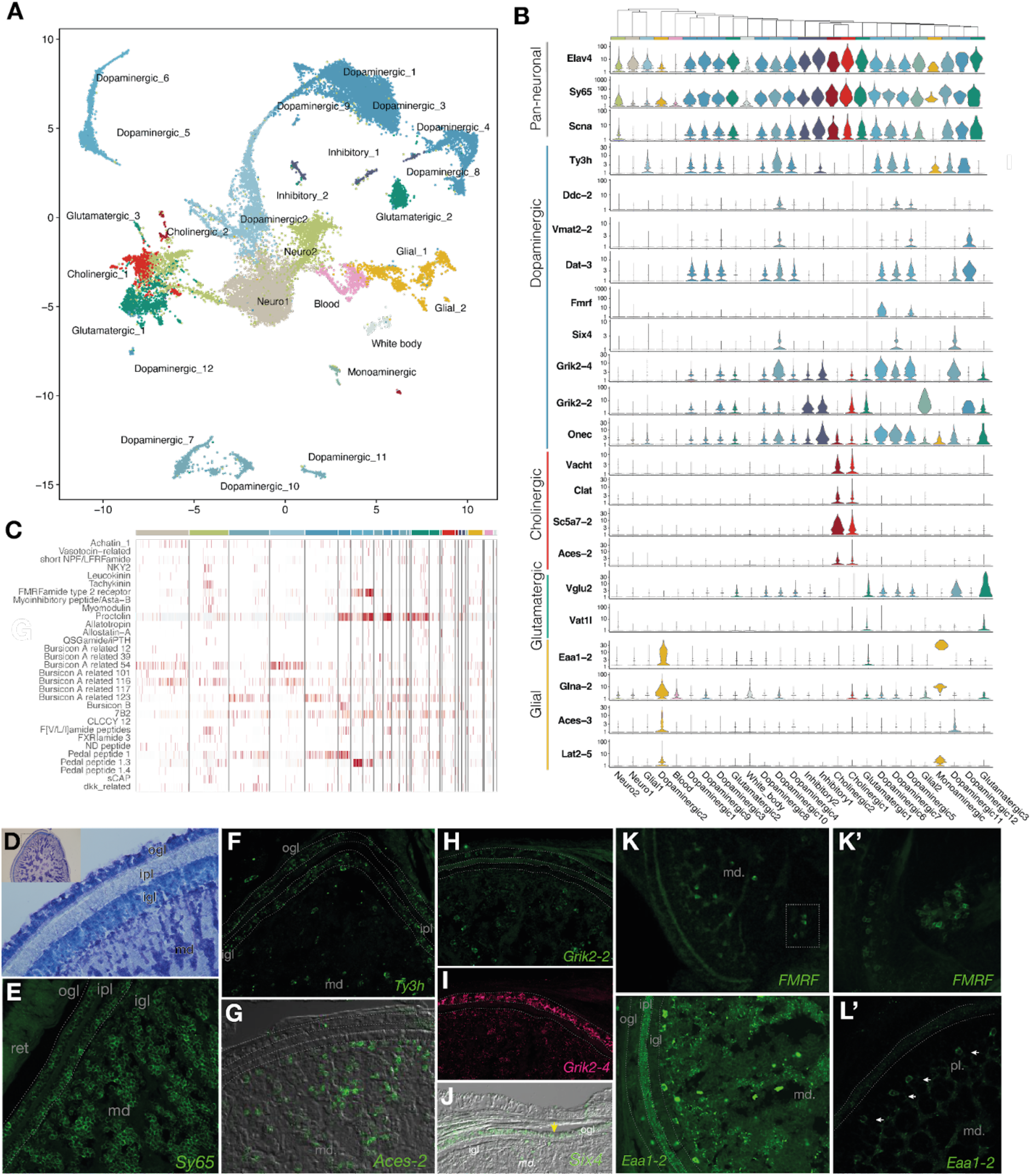
Cell type complement of the *E. berryi* adult optic lobe. (A) UMAP of 26,436 cells from adult *E. berryi* optic lobe highlighting location of each cell population. (B) Violin plots indicating genes specifically expressed in sets of key cell clusters. (C) Heatmap showing expression levels of neuropeptides in each optic lobe cell cluster. (D) Toluidine blue staining of a cross-section of the optic lobe. Inset: overview of optic lobe. (E-L’) Fluorescence imaging of HCR stainings. (E) Pan-neuronal marker Sy65 (synaptotagmin) expression demonstrating location of neuronal cells in both optic lobe cortex and medulla. (F) Dopamine synthesis enzyme Ty3h (tyrosine hydroxylase) expression in a subset of neuronal cells of both optic lobe cortex and medulla. (G) Cholinergic cell marker, acetylcholine esterase, Aces-2, expression in cells of the optic lobe medulla. (H) Kainate glutamate receptor Grik2-2 expression in the optic lobe outer granule layer (ogl) and in large cells of the medulla. (I) Kainate glutamate receptor Grik2-4 expression in optic lobe outer granule layer and punctate expression in the medulla. (J) Six4 expression in the optic lobe outer granule layer of cortex (K-K’) FMRF ligand transcript localization using HCR in wholemounts. (K) Overview of FMRF ligand producing cells arranged in a ‘rosette’ formation located in deep cells of the medulla. (K’) Magnification of FMRF-expressing cell clusters in medulla. (L-L’) Expression of glial marker Eaa1-2 on cryosection (L) and in wholemount staining (L’), demonstrating Eaa1-2 expression in large cells of the medulla including large cells located in palisade layer of the medulla (located directly beneath the inner granule layer of cortex) and a punctate pattern in cells that may be adjacent to projections of cells in inner plexiform layer and of axonal projections in the medulla. Abbreviations: igl - inner granule layer, ipl - inner plexiform layer, md -, ogl - outer granule layer

**Figure 4.**
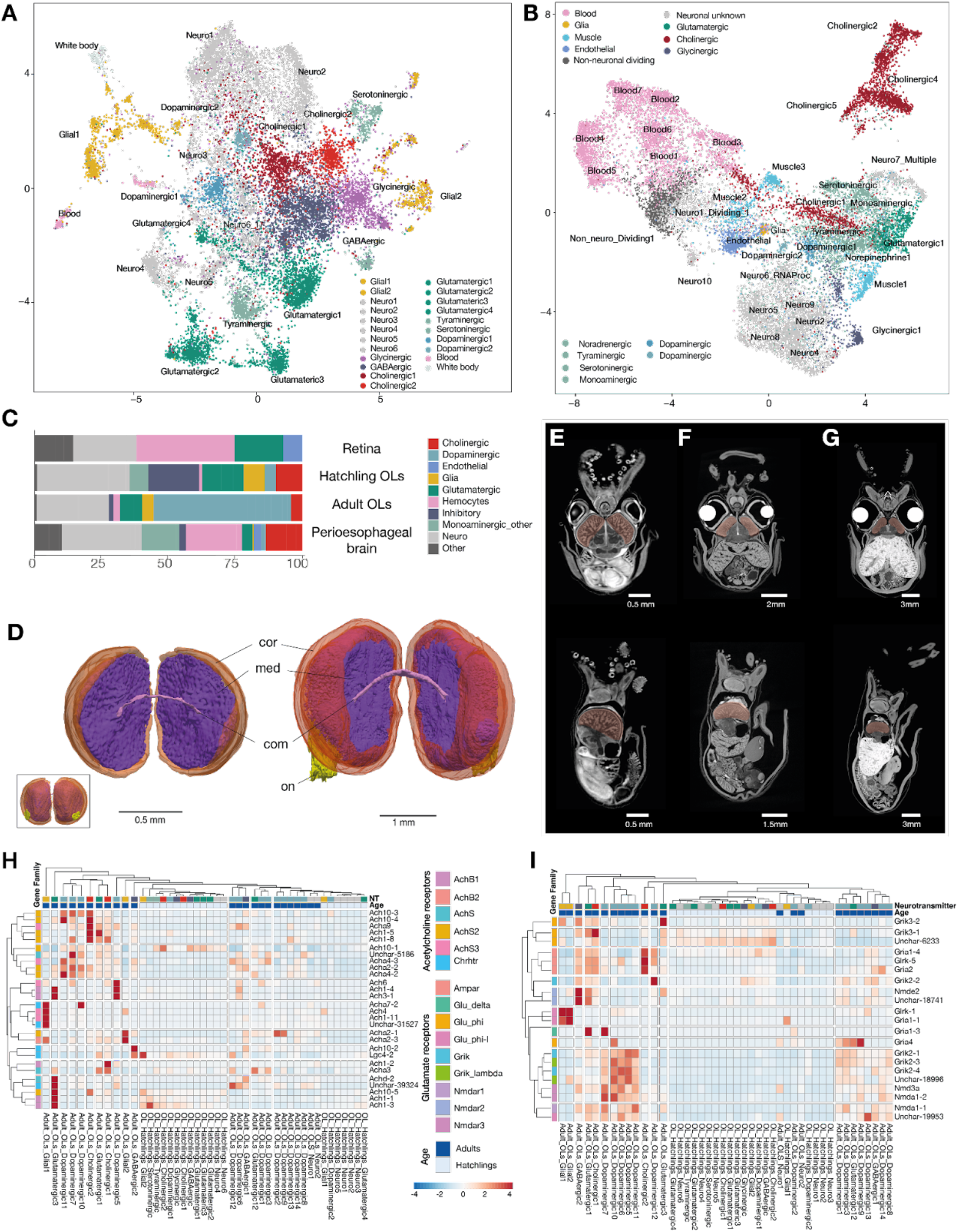
Extended single-cell stage and organ profiling and optic lobe maturation. (A) UMAP of 19,029 cells from 1 dph hatchling. (B) UMAP of 17,468 cells *E. berryi* non-optic lobe perioesophageal brain control dataset. (C) Stacked barplot of number of cells of each cell type in each single-cell dataset. (D) Comparison of optic lobe morphology by microCT segmentation and rendering of optic lobes at 1 day post hatching (left) and 60 days post hatching (right) - posterior view. Cortex is colored in pink with transparency while medulla that lies beneath it is shown in purple (red colored surface corresponds to cortex wrapping the medulla): the bending of the optic lobe causes the cortex to engulf a larger fraction of the optic lobe. Inlet shows anterior view of optic lobes at 1 dph with minute optic nerve located ventrally (yellow). (E-G) MicroCT reconstruction of *E. berryi* over time, at 1 day-post-hatching (E), 40 days-post-hatching (F), 80 days post-hatching (G). The optic lobes are comparatively large and crescent shaped in hatchlings, and become bean shaped in adults (E-G). Transverse view (top) and sagittal view (bottom). (H-I) Heatmaps showing expression levels of cholinergic (H) and glutamatergic (I) markers in adult and hatchling optic lobes. Abbreviations: CNS - central nervous system, com - commissure, cor - cortex, med - medulla, on - optic nerve.

**Figure 5.**
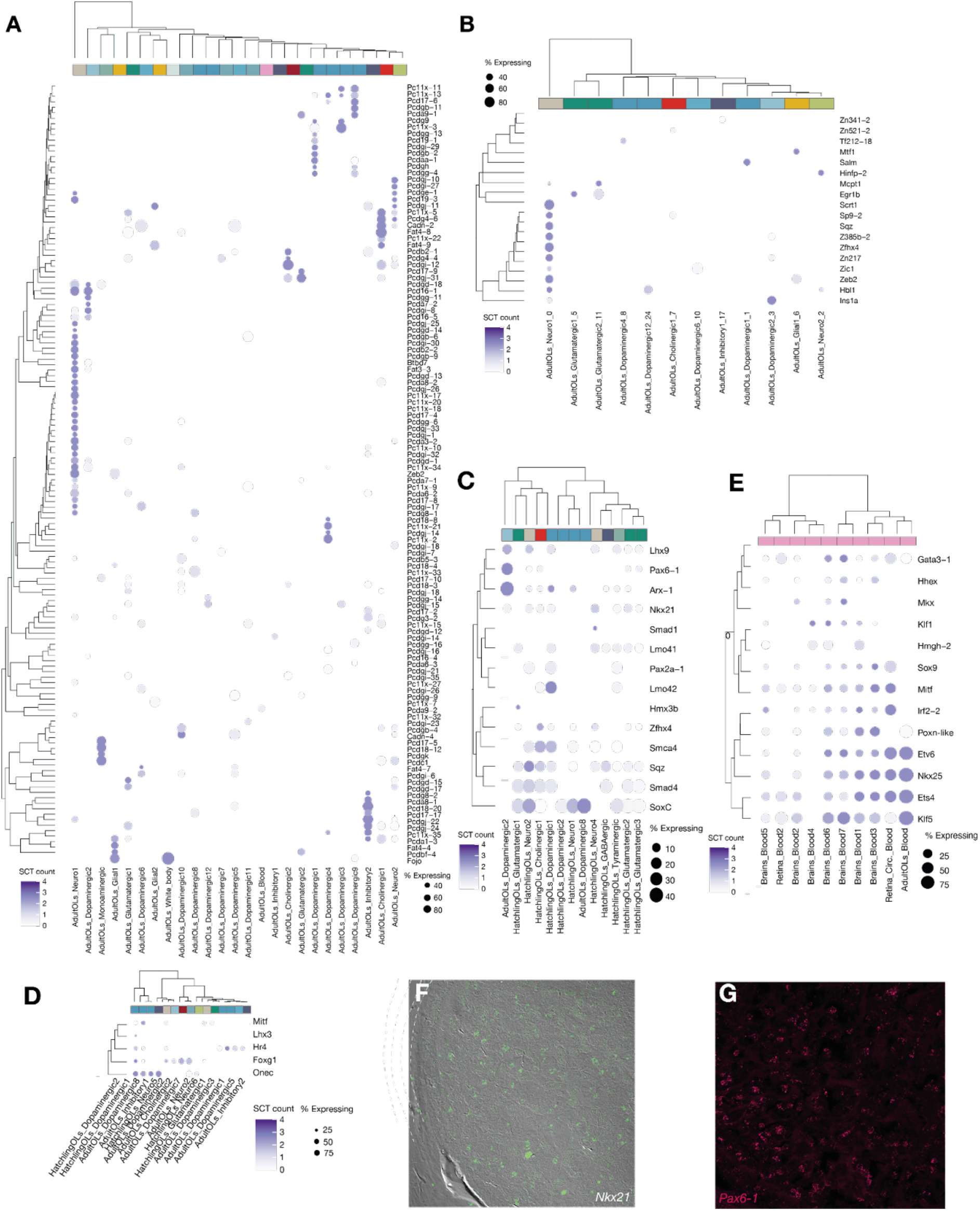
Cell type complexity examined through protocadherin and transcription factor expression. (A) Dotplot showing expression levels of protocadherin molecules in mature adult optic lobes (B) Dotplot showing C2H2 transcription factor expression in mature adult optic lobe. (C-E) Dotplots showing expression of transcription factors in populations of hatchling and mature optic lobes. (F-H) HCR stainings of selected transcription factors (F) Nkx21 expression in the medulla of optic lobe (G) Pax6-1 transcription factor expression in the optic lobe

The eye and optic lobes are highly vascularized, and our dissections necessarily captured the retinal plexus (vasculature underlying the retina), vessels of the optic lobe, and tissues from the white body (a hematopoietic organ situated between the eye and the optic lobe (Figure 1E-F, in white), and the anterior chamber organ, responsible for maintaining fluid in the eye (Figure 1D) (Stephens and Young, 1969; Wild et al., 2015). Accordingly, we found hemocytes in our retina, optic lobe and central nervous system preparations, which we identified by the expression the blood markers Nkx25, Netr-1, and Pgsc2 (Figure 2B, Figure S2E). The nature of these cells was confirmed by scRNA-seq conducted on whole blood from adult *E. berryi* (20,381 cells from four technical replicates). Presumptive white body cells were identified by the co-expression of blood marker Tie1 and Pcna (Castellanos-Martínez et al., 2014; Salazar et al., 2015). These additional control datasets allow us to clearly identify hemocytes in our visual system and brain single-cell analyses.

### The bobtail squid retina displays few neuronal cell types

To characterize the bobtail squid retina, we captured 15,223 cells from dissected adult *E. berryi* retinas (Figure 2A,B). We identified two closely related populations of photoreceptors (Photoreceptor_r1 and _r2) based on their expression of photosensitive molecules, phototransduction cascade effectors (Gq, PLC, Arrestin) and retinol recycling markers (Figure 2C). Of the six putative *E. berryi* photosensitive GPCRs that we identified by phylogenetic analysis – two rhabdomeric opsins (r-opsin1 and 2), two retinochromes (retinochrome 1 and 2) and two xenopsins (xenopsin1 and xenopsin2) (Figure S3B, (Bonadè et al., 2020)) – only the r-opsins and retinochrome-2 showed detectable retinal expression in both scRNA-seq and bulk RNA-seq (Figure S3B). We identify the Photoreceptor_r2 subpopulation as the primary photoreceptor type based on its high expression of r-opsin2, the primary photosensitive molecule in cephalopods (Bonadè et al., 2020). We find that r-opsin2 is also expressed in non-visual tissues in *E. berryi* (Figure S3C, Figure 2C), consistent with r-opsin expression in other cephalopods (Kingston and Cronin, 2016; Kingston et al., 2015a, 2015b).

Unlike Photoreceptor_r2, Photoreceptor_r1 also expresses r-opsin1 and the RABP gene involved in retinol recycling (Figure 2C) (Molina et al., 1992; Terakita et al., 1987; Yau and Hardie, 2009), as well as the neuronal stemness marker Musashi and several neuronal and neurotransmission genes including the vesicular glutamate transporter Vglut and dopamine receptor Dopr1 (Figure 2C, Figure S3). Expression of r-opsin1 mRNA is detected by HCR below the basement membrane suggesting that these cells may occupy a more proximal location in the retina (Figure 2D-D’’). Finally, a small Musashi+ population (‘Photoreceptor_undiff’) also expresses Sy65 and phototransduction markers (but not RABP), as well as an ephrin receptor Epha7 which has been described as a marker of developing photoreceptors in other cephalopod species (Napoli et al., 2021). We hypothesize that Photoreceptor_r1 and Photoreceptor_undiff populations could correspond to different steps in the photoreceptor maturation process.

*E. berryi* photoreceptors express multiple glutamate synthesizing enzymes (*e.g.,* Alat, Got) and transporters (*e.g.,* Glut, VgluT). We did not detect presynaptic markers for acetylcholine, serotonin, dopamine, or octopamine, which have been proposed in the past to play a role in other cephalopod species, or histamine as employed by arthropod photoreceptors (Cohen, 1973; Juorio, 1971; Kime and Messenger, 1990; Stuart, 1999; Suzuki and Tasaki, 1983). Thus, the rhabdomeric photoreceptors of *E. berryi* use glutamate, the same neurotransmitter as vertebrate ciliary photoreceptors, but differ from the histamine used in fly rhabdomeric photoreceptors. Curiously, we observe the expression of innexins Unc7-4 and Unc7-6, that may correspond to components of gap junction-mediated electrical synapses that have been proposed in cephalopods (Yamamoto and Takasu, 1984) (Figure S3D). In a fashion consistent with the ‘deep retina’ hypothesis, we therefore identified a reduced set of glutamatergic photoreceptive neural cells in the retina with populations likely corresponding to successive differentiation steps of photoreceptors.

The squid retina also contains a pigmented cell population (‘Pigmented2’) that is positive for phototransduction genes but negative for neuronal marker Sy65 (Figure S3C). This cell type likely corresponds to the pigmented support cells embedded below the basement membrane of the retina (Koenig et al., 2016; Napoli et al., 2021; Wentworth and Muntz, 1992; Yamamoto and Takasu, 1984; Yamamoto et al., 1965; Young, 1971). These cell types are intermingled in the retina as schematized in Figure 2H and can be readily identified on corresponding EM sections (Figure 2H). A second pigmented population (‘Pigmented_tyro1’) features the tyrosinase (Tyro) gene that is expressed in a ring-like domain around the aperture of the pupil (Figure 2G).

In bobtail squid, the lens is secreted by cells at the periphery of the retina. The lens of coleoid cephalopods comprises two classes of crystallins of independent phylogenetic origins: cephalopod-specific Glutathione-S-Transferase-derived lens proteins (σ-crystallins) (Koenig et al., 2016; Tan et al., 2016; West et al., 1994) and spiralian-specific aldehyde dehydrogenase-like proteins (Ω-crystallins) (Belcaid et al., 2019; Piatigorsky et al., 2000). We found a cluster of lentigenic cells expressing both Ω-crystallin and σ-crystallins (Arnold, 1967; Tomarev and Piatigorsky, 1996; West et al., 1994). HCR staining confirmed that both crystallin classes are co-expressed in lentigenic cells that surround the lens (Figure 2F-F’). Thus, the cephalopod lens curiously incorporates elements of exaptation of multiple lineage-specific proteins (based on their useful refractive index). Moreover, lentigenic cells also expressed Sp9, a member of the Krüppel-like class of transcription factors involved in *D. pealei* lentigenic cell specification (Figure 5) (McCulloch and Koenig, 2020).

### The optic lobe utilizes a complex cell type catalog

The photoreceptor axons leave the squid retina in bundles surrounding the eye and enter the optic lobes latero-ventrally. The squid optic lobes are composed of a cortex (Ramón y Cajal’s ‘retina profunda’) and a medulla (Fig.1 D-E, Figure 3D (Dilly et al., 1963)). The cortex comprises two granule layers containing neuronal cell bodies and a plexiform layer containing neurites and no cell bodies, as shown by Sy65 expression (Figure 3E). In other cephalopods, photoreceptor axons have been described to run through the outer granule layer without forming synapses and then synapse with elements of the plexiform layer or the outer part of the medulla, the palisade layer (Dilly et al., 1963; Young, 1974) (Figure 1AE-G, Figure 3D-L’).

To characterize the cell type diversity of the optic lobe, we profiled 26,436 cells from mature adult optic lobes (>60 days post hatching, Figure S2A-D), and identified 26 major cell populations in the optic lobe after dimensionality reduction (Figure 3A), of which 22 expressed the pan-neural marker Elav4. Twenty Elav4+ correspond to differentiated neuronal cell populations (Elav4+, Sy65+ and Scna+), two clusters (Neuro_1 and Neuro_2) constitute undifferentiated neuronal cells (Elva4+ but Sy65-), and four clusters represent non-neuronal types (Figure 3A-B). The majority of neuronal cell clusters in the optic lobe were dopaminergic (12/22) expressing both dopamine synthesizing enzymes (Ty3h, Ddc-2) and transporters (Dat-2, Vmat2-2) (Figure 3A-B). We also identified two cholinergic cell clusters (expressing the acetylcholine synthesizing enzyme ChAT, acetylcholine transporter VAchT, choline transporter Sc5a7-2 and acetylcholine degrading enzyme Aces-2); three glutamatergic cell types (featuring vesicular glutamate transporter VgluT and Vat1l, Figure 3A-B), and two putative inhibitory cell clusters expressing some GABAergic markers (GAT, Gabt-1, Gabp2, Figure 3A). Low levels of ChAT were also detected in a few cells of cluster Neuro_1. Additionally, we observed a small cluster expressing Vmat2-2 that was considered monoaminergic. Each neuronal cluster expresses a distinct complement of neurotransmitter receptors, implying that these neurons receive diverse inputs, including a cluster-specific set of acetylcholine and monoaminergic neurotransmitter receptors (Figure S4A-B). Glutamatergic cluster 1 expresses the largest number of monoaminergic receptors (Figure S4A). Curiously, dopaminergic neurons also expressed low levels of VgluT, the primary vesicular glutamate transporter in the *E. berryi* genome (Figure 1J). The presence of VgluT in dopamine-synthesizing cells is reminiscent of observations of glutamate-dopamine co-transmission observed in vertebrate species both during embryonic development and in adult neuronal structures that may play a role in response to toxins (Eskenazi et al., 2021).

### A retrograde signal involving dopamine and FMRF signaling

Distinct types of dopaminergic (i.e., Ty3h+) neurons were found by HCR in both the cortex and the medulla of the optic lobe (Figure 3F). We assigned three dopaminergic populations (Dopaminergic 7, 10, 11; Figure 3A-B) to the outer granule layer of the cortex based on the expression of the Six4 transcription factor and kainate glutamate receptor Grik2-4 (Figure S3B,J). Other dopaminergic populations (Dopa5,6 and Dopa1,3,9; see below) appear to be located in the medulla (Grik2-2+, Figure 3H). Cholinergic (Aces-2+) cells were observed in various locations within the medulla including the palisade layer situated directly beneath the cortex (Figure 3A-B,G). Notably, some phylogenetically closely-related genes exhibit distinct expression domains. For instance, the paralogous kainate glutamate receptors Grik2-2 and Grik2-4 were observed in the outer granule layer and in large cells of the medulla with complementary expression patterns and in distinct cell clusters (Grik2-2 in clusters 1, 3, 9 in and Grik 2-4 in the dopaminergic cells of the cortex (Dopaminergic 7, 10, 11) and centrifugal (Dopaminergic 5,6 clusters, see below (Figure 3B,H-I)).

Among the arguments in favor of Ramón y Cajal’s ‘deep retina’ hypothesis relating the cortex of the cephalopod optic lobe to the vertebrate neural retina was the observation that some cells in the cephalopod optic lobe medulla project back in the direction of the retina (Young, 1962a, 1974). We identified FMRF-amide as a potential retrograde signal from optic lobes to the retina in *E. berryi*. FMRF peptide was detected in the retina through antibody staining (Figure S5A), but its encoding transcript was not found in the retina through scRNA-seq, bulk RNA-seq or HCR. We did, however, recover prominent expression of both transcript and peptide in the optic lobes (Figure 3A,C). Dopaminergic populations 5 and 6 (top left of the UMAP view, Figure 3A), produce FMRF-amide transcript and were observed using HCR in the cells of the optic lobe medulla, organized in a rosette formation (Figure 3C,K-K’). These cells match the features of the ‘centrifugal’ cells described by J.Z. Young (Young, 1962a) and is consistent with the suggestion of retrograde projections from the medulla to the retina (Ramón y Cajal, 1930; Young, 1962a, 1974). This finding is consistent with recent whole-cell patch-clamp recordings from centrifugal neurons of the optic lobe showing reactivity to FMRF-amide (Chrachri, 2020). These may be functionally analogous to the lateral inhibition of photoreceptors mediated by horizontal cells in the vertebrate retina, which relies on distinct signaling mechanisms (Kramer and Davenport, 2015).

### Identification of glial cell population

Glial cells are non-neuronal support cells that are present in most complex nervous systems, but whose evolutionary origin and homology remain unclear (Hartline, 2011). We identified several putative glial cell types in *E. berryi* among non-neuronal (i.e., Elav4-, Sy65- and Scna-) cell populations. Although there are no known glial markers in cephalopods (Ibrahim et al., 2020; Imperadore et al., 2017), the glial identity of these cells is supported by the absence of any presynaptic markers and presence of neurotransmitter degrading enzymes (Glna-2 and Aces-3) and transporters (Lat2-5 and an Eaat), which have, in the case of Eaat transporters, been associated with both glial and presynaptic cells in vertebrates (Rothstein et al., 1994). Our analysis provides a first catalog of marker genes enriched in the glial cells of *E. berryi* (Figure S2F, Table S.6-9). Notably, we did not observe any expression of orthologs of arthropod glial markers such as Repo and Gcm, or evidence of presence of vertebrate markers such as GFAP or S100 in the *E. berryi* or in other cephalopod genomes (Albertin et al., 2015b; Belcaid et al., 2019), which supports the convergent evolution of glial cells across diverse bilaterians (Hartline, 2011).

The glial marker Eaat1-2 was expressed in both large cells situated in the palisade layer of the medulla (Figure 3J-J’, arrowheads) and in a punctate pattern in the inner plexiform layer and was also associated with putative neurites of the medulla reminiscent of the putative glial cells observed in *Octopus* (Dilly et al., 1963). Interestingly, Eaat mRNAs have recently been reported to localize to endosomes in vertebrates (Popovic et al., 2020). These two distinct patterns may correspond to the two distinct putative glial clusters in *E. berryi* optic lobes (‘Glial_1’ and ‘Glial_2’)(Figure 3A-B,L-L’).

### Cell types diversity in hatchlings suggests optic lobe maturation

The eyes and optic lobes of cephalopods grow considerably over the life of the animal (Liu et al., 2017) and underlie a range of age-specific visual behaviors. For example, one day-old *E. berryi* hatchlings exhibit light sensitive behaviors such as burying and visually-driven hunting and catching of prey, implying that their visual system is functional (Figure S5B, Supp. Data video1 & 2) as observed in other cephalopod species (Guibé et al., 2012). Unlike hatchlings, however, mature animals also exhibit additional visual behaviors including those relating to mating (Gutnick et al., 2021). To begin to characterize changes in the visual system with age, we characterized the morphological differences in the optic lobes of *E. berryi* using microCT. We observed changes in the shape of the optic lobes (from half-sphere to kidney bean-shape, Figure 5E-H) throughout their post-hatching life, but also a change in the position and size of the optic nerve and commissural neurons. Notably, the cortex thickness increased between 1-80 dph and that the ratio of the cortical and medullar volumes remained constant, indicating isometric growth of these two closely integrated structures (Figure 1F, Figure 4E-G, Figure S5B, Table 2).

To study the impact of the growth of the optic lobe from hatchling to adult on cell types, we profiled 19,029 cells from the optic lobes of one-day old *E. berryi* hatchlings, and compared them with the adult samples described above (Figure 1C and Figure 4A). Glial, hemocyte and white body populations expressed similar markers to those observed in the adult optic lobes (Figure S2E-F). Surprisingly, however, the complement of neuronal cell types we recovered differed substantially between hatchling and adult, notably accompanied by an increase in the number of dopaminergic cells and cell types (Figure 4A,C). Specifically, we recovered fewer dopaminergic cells in hatchlings than in the mature optic lobe, with only two dopaminergic clusters, along with four glutamatergic, two cholinergic and six neuronal populations that could not be assigned to a specific neurotransmitter (Figure 4A,C). The combination of neurotransmitter receptors observed in the mature optic lobes is different from the one in the hatchling optic lobe samples (Figure S4A-B, Figure 4H-I). In contrast to adult optic lobes, we also observe tyraminergic and serotonergic cell types in hatchling optic lobes that were not captured in discernible numbers in the adult optic lobes scRNA-seq, as well as an additional small population of glycinergic inhibitory neurons.

To test whether increased usage of dopamine is a general hallmark of central nervous system maturation or a particularity of the optic lobes, we analyzed an additional 17,468 cells from the peri-oesophageal brain lobes (i.e., non-optic lobe brain) from the same mature adult individuals. We found a complex set of neuronal cell types (Figure 4B). Unlike the optic lobe, however, dopaminergic cells were not predominant. While this control dataset is insufficient to fully characterize the full cellular complexity of the cephalopod brain, it extends our classification of cell-types based on neurotransmitter type beyond the optic lobe.

### From PCDH to neurotransmitter receptors: molecular contributions to cell type diversity

We further explored the molecular contributors to cell type diversity by examining cell adhesion, neurotransmitter receptor, and transcription factor genes associated with nervous system complexity in other lineages (Figure 1J). The broad expansion of protocadherins (Pcdh) and C2H2 zinc finger transcription factors (C2H2s) in cephalopods was predicted to play a key role in specifying their neuronal repertoire (Albertin et al., 2015b). In *E. berryi*, we found 320 genes bearing Protocadherin domains (Figure 5A-B, Figure S5C-F). Indeed, we observed protocadherin expression predominantly in neuronal tissues including photoreceptor cells of the retina consistent with their role in neuronal specification (Pancho et al., 2020), but also in endothelial and immune cells suggesting possibly additional roles (Dilling et al., 2017) (Figure 5A, Figure S5B, G-H). Although most expressed PCDHs belonged to the expansions on chromosome 20 (ALGs I and G), others expressed in dopaminergic (1 in hatchling and adults) and glial cluster, as well as photoreceptor_r1 in the retina were encoded on other chromosomes. While some cell clusters expressed extensive repertoires of PCDHs, consistent with a role as interneurons, others, such as the adult optic lobe dopaminergic 5 and 6 or retinal photoreceptor_r2 clusters, did not express PCDHs (Figure 5A, Figure S5B,G-H). Similarly, the optic lobes of hatchlings expressed a wider variety of overlapping sets of C2H2s (Figure S5D-F), whose expression decreases considerably in mature optic lobes (Figure 5B).

The expression of cholinergic receptors in multiple cell type clusters (Figure S4A) indicates that cholinergic neurotransmission plays a more widespread role than monoaminergic neurotransmission in the cephalopod optic lobes (Figure S4B). Moreover, a specific ‘code’ of metabotropic receptors and ionotropic receptor subunits seemingly designate different cells that receive cholinergic signals (Figure S4B). Often, both spiralian and bilaterian families were expressed in the same cell clusters suggesting they could take part in hetero-pentamers. Moreover, the cholinergic and glutamatergic receptor codes (Figs. 4I and 4J, respectively) were more prominent in adults than in hatchlings, consistent with an overall maturation of the optic lobe.

Transcription factors have been proposed as determinants of cell type identity conserved across lineages (Arendt et al., 2016). We expected to observe canonical retinal determination genes in our retina dataset (homologs of atonal, sine oculis, dachshund, eyes absent and Pax6) (Martín-Durán et al., 2012). However, possibly because these genes are associated with embryonic development (Yoshida et al., 2014) and cell fate specification rather than with adult cells, we did not observe strong co-expression of these genes in our hatchling or adult samples. The bilaterian master eye regulator Pax6 (Chow et al., 1999; Gehring, 2005) is only subtly expressed in adult *E. berryi* photoreceptor populations, and prominently in the contractile cells surrounding the eye. In the optic lobe, it is present in the putative centrifugal-like cells of adult optic lobe together with Egl13 (dopaminergic clusters 5 and 6) and in an additional dopaminergic population (Dop 2, Figure 5C, H). A sine oculis ortholog (Six4) is a marker for three dopaminergic populations (7,10 and 11), and is observed together with Grik2-2 in the cortex of the optic lobe in cells that may correspond to the amacrine cells described by J. Z. Young (Young, 1962a, 1974). Nkx21 was expressed in cholinergic cell types in both mature optic lobes and perioesophageal brain (Figure S4C, Figure 5C). Similarly, both Aces-2 and Nkx21 mRNAs are localized in the medulla (Figure 3I, Figure 5H), we suspect that these cholinergic cells may correspond to the bipolar cells of the medulla described by J. Z. Young (Young, 1962a, 1974). Glial cells were specified by a combination of transcription factors including Foxg1 in mature tissues, Hr4 in hatchlings, and Sun and Foxp1 in mature optic lobes, possibly indicating a larger diversity of glial cell fates in cephalopods that previously recognised (Stephens and Young, 1969). In contrast, hemocytes across samples expressed known mesodermal factors including Klf5, Ets4, Nkx25, Etv6, Sox9 and Gata3-1 (Figure 5E,G, Figure S4C). Ets4 expression has been reported in the cortical layers of embryonic optic lobes of *Idiosepius (Yoshida and Ogura, 2011)*, but was not observed in the hatchling and adult *E. berryi* optic lobe cortex.

Overall, while we found that PCDH and various neurotransmitter receptors play a major role establishing the diversity of cell types in bobtail squid, we identified only limited conservation of transcription factors expression between cephalopods and fly or vertebrates.

### The contribution of gene evolutionary history to organ-level convergence

Although the bobtail squid visual system has a very different anatomical organization than found in vertebrates and *Drosophila* (Figure 1A, Figure 2), we nevertheless investigated possible cell-type homologies between *E. berryi* and other bilaterians. To compare homologous gene usage across species, we used gene family reconstruction to pinpoint the origin of the genes that define cell type identity (as computed using a Wilcoxon rank sum test, Tables S6-9). In both *E. berryi* retina and optic lobe we found that most of the markers defining cell types belong to pan-bilaterian gene families (Figure 6A-B, Figure S6A-B). In general, non-neuronal cell types such as blood, white body or glial cells, as well as putative progenitor cell populations (e.g., Neuro1 & 2 in adult optic lobes) are designated by markers of more recent phylogenetic origin than differentiated neuronal cell types, with the exception of retinal photoreceptors that possess cephalopod-specific markers (Fisher enrichment test, Figure S6A).

**Figure 6.**
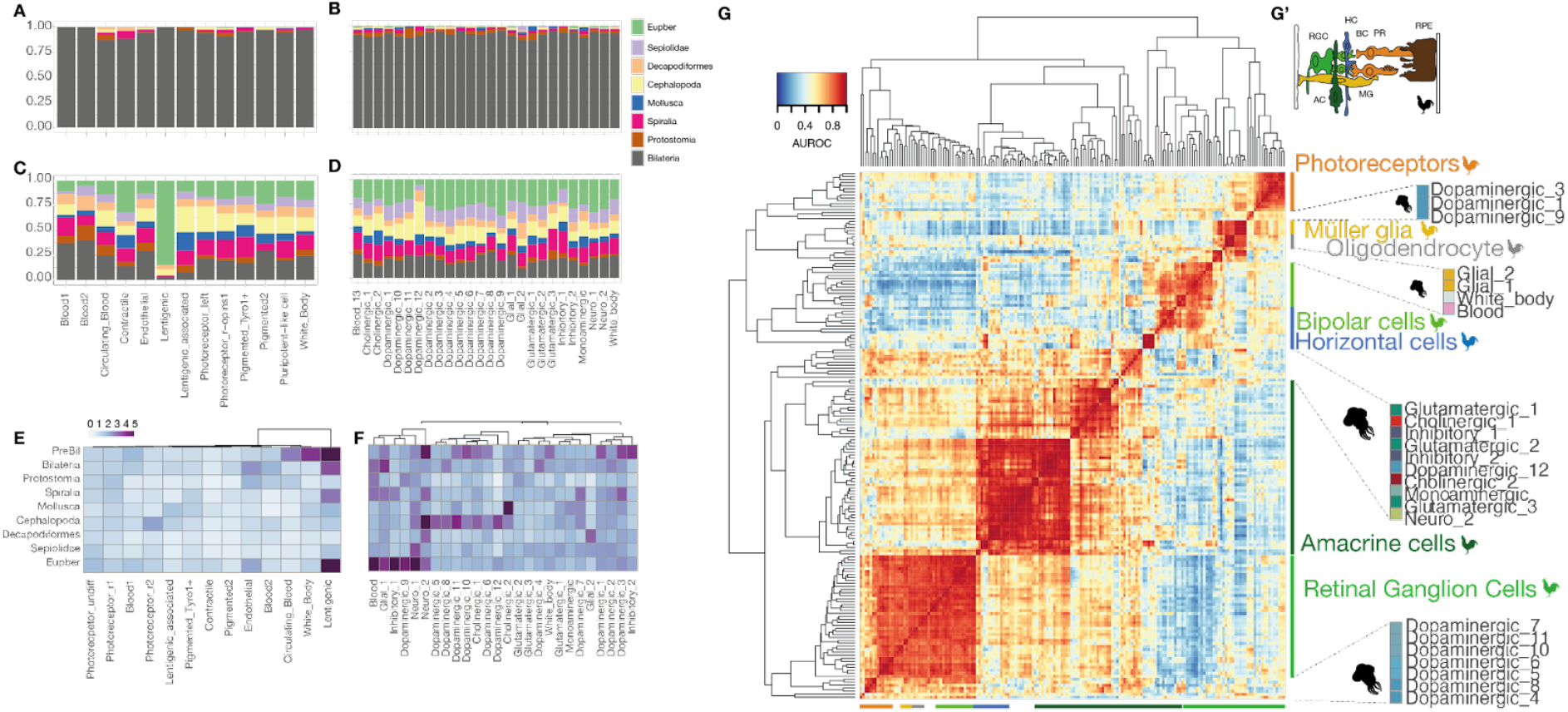
Evolutionary history of genes employed by *E. berryi visual* system. (A,B) Stacked barplots indicating the fraction of genes originating from gene families at each strata in (A) retinal and (B) adult optic lobe cell populations (C,D) Stacked barplots indicating the fraction of genes whose last gene duplication dates to each strata in (C) retinal and (D) adult optic lobe cell populations. (E,F) Heatmaps displaying significance of enrichment for a given phylostrata (-log10(p-value) of Fisher’s exact for the enrichment of genes whose last duplication took place in each strata (E) retina and (F) adult optic lobe cell populations. (G) Heatmap displaying AUROC scores or “Reciprocal_top_hit’’ match types are identified as similar cell-type pairs in comparisons between *E. berryi* adult optic lobe cell populations compared to cell types of the chicken retina (Yamagata et al., 2021), (G’) inset showing location of cell types in vertebrate retina (reproduced from (Brady et al., 2005).

To gain further insight into the evolutionary history of genes, we traced the duplication events that give rise to current genes using a gene-species tree reconciliation approach (Morel et al., 2020). We determine that while many gene families have ancient origins, most (2342/3759, 62.3%) cell-type specific genes have undergone more recent duplication events. This observation is consistent with our description of duplication and gene losses in the neural gene repertoire (Figure 1J). Notably, in the optic lobe, seven cell types (cholinergic and dopaminergic) are enriched in genes duplicated at the cephalopod node, while five other cell populations, including glia, blood and inhibitory or undifferentiated neurons are enriched in marker genes duplicated more recently in *E. berryi*. For instance, the cholinergic cluster 1 features potassium channels (Kcna-1 and Kcnn-1) and acetylcholine receptors (Ach1-3, Acha4-3) that are duplicated in cephalopods (Tables S11-12). In the retina we observe somewhat fewer expressed lineage-specific gene duplications, except in the lentigenic cells that are characterized by the expression of duplicated σ-crystallins. These findings emphasize the fact that despite pan-bilaterian conservation of neurotransmitters and their synthesis enzymes, the genes involved in neuronal cell types have a complex evolutionary history that participated in independent origins of many neural cell types.

To compare cell types across species, we measured similarity of expression patterns between homologous genes using a pairwise unsupervised analysis (MetaNeighbor) (Crow et al., 2018; Wang et al., 2021). We compared the cell types of the bobtail squid optic lobe with those of the chicken retina (Yamagata et al., 2021) (Figure 6G), the adult fruit fly optic lobe (Özel et al., 2021) (Figure S6E) as well as those of mouse brain regions involved in visual processing, such as the dorsal lateral geniculate cortex (Allen Mouse Brain Atlas [LGN], 2011), Figure S6D) and visual cortex (Tasic et al., 2016), Figure S6E). We also assessed possible cellular correspondences between the retinas of bobtail squid and chicken (Figure S6A). Neuronal cell types are primarily grouped together by species and according to their neurotransmitter in the similarity-based clustering, and few groupings gather cell types from both species. Some excitatory neuron types appear similar across species, particularly some dopaminergic cell types (1,3 and 4-11) of the squid optic lobe that resemble mouse glutamatergic cells of the visual cortex or the retinal ganglion cells (Figure 6G and S5E). Interestingly, the retinal ganglion cells show some affinity for bobtail squid photoreceptor, which can be explained by their shared usage of rhabdomeric melanopsin and associated transduction pathways (Isoldi et al., 2005). Moreover, inhibitory cell types appear less similar to each other across species than excitatory ones (e.g.. OL glutamatergic cells).

As expected, we observed a strong separation between neuronal and non-neuronal cell types. In contrast, glial cell types in bobtail, chicken or mouse show higher similarity than neurons, but also notably formed a group with other cell types related to blood, white body and endothelium. This observation suggests that increased similarity scores in these cell types that are ‘outgroups’ to the neuronal ones are caused by the weak similarity between neuronal and non-neuronal cells rather than by evolutionary conservation. Given the biological and functional disparity of these cell types, their apparent similarity can be explained by a ‘long branch attraction’, consistent with the consensus on the convergent evolution of the glia in cephalopods and vertebrates (Hartline, 2011). Altogether, our cross-comparisons argue for a reduced repertoire of ancestral neuronal cell types in bilaterians, as they support the lineage-specific elaboration of each neurotransmitter-specific cell population. They also do not suggest a direct correspondence between the cell types of the cephalopod optic lobe cell types and the vertebrate retina, rejecting a literal interpretation of Ramón y Cajal’s “deep retina” hypothesis.

## Discussion

We used comparative genomics and single cell analysis to investigate the convergent visual systems of cephalopods and vertebrates, both of which are uniquely endowed with camera-type eyes (Kozmik et al., 2008). The cephalopod visual system therefore constitutes a unique model for understanding the underpinning of this convergent evolution at the molecular, cellular and organ levels. The annotated genome of the bobtail squid *E. berryi* provides a platform for detailed studies in this emerging model. To go beyond previous organ-level transcriptomic studies (Ogura et al., 2004; Yoshida and Ogura, 2011), we set out to investigate gene expression in cell types and in doing so evaluate the ‘deep retina’ hypothesis formulated by Ramón y Cajal (Ramón y Cajal, 1930).

The cephalopod retina has fewer cell types than the complex vertebrate retina, suggesting that visual signal processing occurs in the cephalopod optic lobe. The 26 cell types we find in the bobtail squid optic lobe cannot be placed in simple correspondence to the broad cell classes (amacrine, horizontal, bipolar, etc.) described in vertebrate retinas. As with vertebrates, we anticipate that more extensive studies will subdivide these 26 cell types into subtypes based on function, connectivity and expression of marker genes (Cheng et al., 2021; Shekhar et al., 2016, 2021; Tran et al., 2019). In vertebrates, the ability to drive reporter gene expression and enrich for specific cell classes by cell sorting has been essential for detailed analysis; these tools are not yet available in cephalopods.

The ‘deep retina’ hypothesis implies an absence of visual signal processing in the retina, the presence of various neuronal cell types dedicated to signal processing in the optical lobe, including retrograde projections from optic lobe to retina (Ramón y Cajal, 1930; Young, 1974). We indeed did not find interneurons within the *E. berryi* retina, neither detectable in the TEM (Figure 2H) nor by scRNA-seq (Figure 2A), in line with observations by J.Z. Young (Young, 1960, 1962b). In the optic lobe,we found three related dopaminergic cell types playing a putative interneuron role that we localized to cortical layers that Ramón y Cajal called the ‘deep retina’. Their joint expression of a repertoire of protocadherin molecules and of several glutamate receptors (Figure 4H) indicates a role in integrating glutamatergic input from the retina. We also find two dopaminergic cell types, located within the medulla, that likely correspond to the ‘centrifugal cells’ mediating retrograde signals described by J.Z. Young (Young, 1962b). These cells express the Pax6 transcription factor and FMRF-amide transcript; the FMRF-amide peptide, but not its transcript, can be detected in the retina (Figure S4C, Figure 5). Our finding is also consistent with recent findings of FMRF sensitivity of centrifugal neurons in cuttlefish (Chrachri, 2020). In vertebrates, lateral inhibition of photoreceptors is mediated by retrograde signals from horizontal cells, although the specific signals are not known (Kramer and Davenport, 2015)

Another cell population described outside the ‘deep retina’ in earlier studies were the bipolar cells of the medulla, which may correspond to the cholinergic cell clusters expressing Nkx21 and Aces-2 found in our study (Figure 3B,G, Figure 5H). The extensive expression of cholinergic receptors throughout the optic lobe (Figure 4 I, Figure S4B) further points to the importance of acetylcholine in optic lobe function. We have therefore identified populations that play presumptive input and output roles, and possibly interneurons interacting through adhesion molecules such as PCDHs, laying the foundation for reconstructing the circuitry of the bobtail squid visual system (Sanes and Zipursky, 2010). Further molecular characterization of the interaction affinity of adhesion molecules in cephalopods will help achieve this goal, for instance by determining which PCDHs mediate homo- or heterophilic interactions (Honig and Shapiro, 2020).

We observed several similarities to vertebrates and *Drosophila* that have not previously been reported in cephalopods. First, we observed two populations of photoreceptors, one of which expressed a second r-opsin gene. Although two r-opsin genes have been described in cephalopods, expression of more than one r-opsin in the retina has not been described (Bonadè et al., 2020). The presence of two photoreceptor populations with multiple opsin molecules is curious as cephalopods are not known to rely on color vision, but rather to detect differences in the polarization of light (Marshall and Messenger, 1996; Mäthger et al., 2006; Messenger et al., 1973; Saidel et al., 2005). Moreover, we identify glutamate as the primary neurotransmitter used by photoreceptors, based on their expression of glutamatergic presynaptic vesicular transporter Vglut (Figure S3A,C). The usage of multiple different neurotransmitter in different cephalopod species has been proposed (Bacq, 1935; Cohen, 1973; Florey and Florey, 1954; Juorio, 1971; Kime and Messenger, 1990; Lam et al., 1974; Loe and Florey, 1966; Makman et al., 1987; Suzuki and Tasaki, 1983). At present, we cannot confirm whether this is due to interspecific differences. The use of glutamate by the rhabdomeric photoreceptors of *E. berryi* which is the same neurotransmitter as vertebrate ciliary photoreceptors, rather than histamine used in fly rhabdomeric photoreceptors further highlights the unique parallel evolution of cephalopods.

At the gene level, we found that while the genes establishing basic neuronal function such as neurotransmitter synthesis (e.g., Ty3h, ChAT) were conserved in limited copy numbers in bilaterians, an extensive diversification of molecules such as receptors, channels or adhesion molecules (e.g., PCDHs) likely played a key role in the evolution of new lineage-specific neuronal cell types (Figure 1I, Figure 6B,D). At the cell type level, this molecular diversification makes it difficult to compare gene expression across species, especially at large evolutionary distances (see Methods). Transcriptomic similarity across neuronal cell types of bobtail squid, chicken, mouse or Drosophila does not reveal obvious deep homology for most neuronal cell types, suggesting a strong lineage-specific drift in gene expression for neuronal cell types as seen in other evolutionarily deep comparisons of neurons (Cardoso-Moreira et al., 2019; Sebé-Pedrós et al., 2016). This is corroborated by the distinct sets of transcription factors that specify neuronal identities in the bobtail squid compared to insect or vertebrate species. We observe few cross-specific associations of neuronal cell types that always involve excitatory neuronal types but disregard other neurotransmitter types (Figure 6G). This suggests that inhibitory neurons might be more prone to convergent evolution and that the bilaterian ancestor likely had several distinct neuronal cell types from which lineages specific complexification of nervous systems took place (Martín-Durán et al., 2018). Interestingly, comparison between reptiles and mammals indicated that excitatory neuronal types are more divergent whereas inhibitory types are more conserved (Tosches et al., 2018). The evolutionary distances separating mammals and reptiles are however comparable to those separating different cephalopods, thus highlighting differences in conservation patterns at distinct evolutionary scales. Similarly, the glial cell populations identified in *E. berryi* are not obviously related to glial cell types found in other species, and do not express the typical transcription factors found in arthropod and vertebrate glia (Hartline, 2011). An investigation of the cell types in the visual systems of other molluscs, such as the sea scallops that possess mirror eyes, could shed light on their evolutionary trajectories.

Furthermore, we observed a change in morphology and striking differences in cell type complements of optic lobes between hatching and the first few months. *E. berryi* are active at night and spend the day buried under the sand (Sasaki, 1929). Unlike the Japanese flying squid *Todarodes pacificus* whose front appendages are fused together at hatching (Watanabe et al., 1996) or other cephalopods that undergo a paralarval stage (Young, 1988), *E. berryi* hatchlings are capable of hunting and display light-evoked behaviors (Figure S5B, Video S1, S2). However, their swimming and predatory behavior becomes more refined with age (Darmaillacq et al., 2014; Villanueva et al., 1997), which may be reflected in the possible maturation process that we observe. Interestingly, two recent preprints (Duruz et al., 2022; Styfhals et al., 2022) describe the cell types in the head of hatchling and embryos of two distinct cephalopod species, and their cell clustering, show similarities with what we observe in the optic lobe at the hatchling stage. Such a change in cell type complement is reminiscent of the situation reported in the *Drosophila* ‘optic lobe’ which is not related either evolutionarily or embryologically to the cephalopod optic lobes (Ma et al., 2012). There, several neuronal clusters die prior to adulthood, but neuronal cell populations that are indistinguishable at the transcriptome level during early stages of neuronal development go on to yield cells with distinct gene expression profiles, morphologies and connectivity (Özel et al., 2021).

Cephalopods are renowned for their cognitive capabilities, learning and social structures (Gutnick et al., 2021). Our single-cell datasets in the genomically-enabled *E. berryi* reveals an exquisite diversity of cell-types, which show distinct distribution in the optic lobe and brain, but also appear to fluctuate during post-hatching, consistent with the existence of a possible maturation process. The existence of adult plasticity is the hallmark of complex nervous systems such as those of vertebrates. The heavily rearranged genome architecture of cephalopods with the fusion of multiple ancestral linkage groups might have not only promoted duplication hotspots, but also permitted the acquisition of more complex gene regulatory processes that could have promoted a cell type specific expression of duplicated genes as seen, for example, in vertebrates (Albertin et al., 2022a; Marlétaz et al., 2018). We hope that our single-cell atlas coupled with the model capabilities of the *E. berryi* will promote further studies to decipher the underpinnings and evolutionary origin of cephalopod nervous systems and detect distinct subtypes as has been the case for *Drosophila* and vertebrate models (Konstantinides et al., 2022; Sanes and Zipursky, 2020; Shekhar and Sanes, 2021).

---

**Figure S1.**
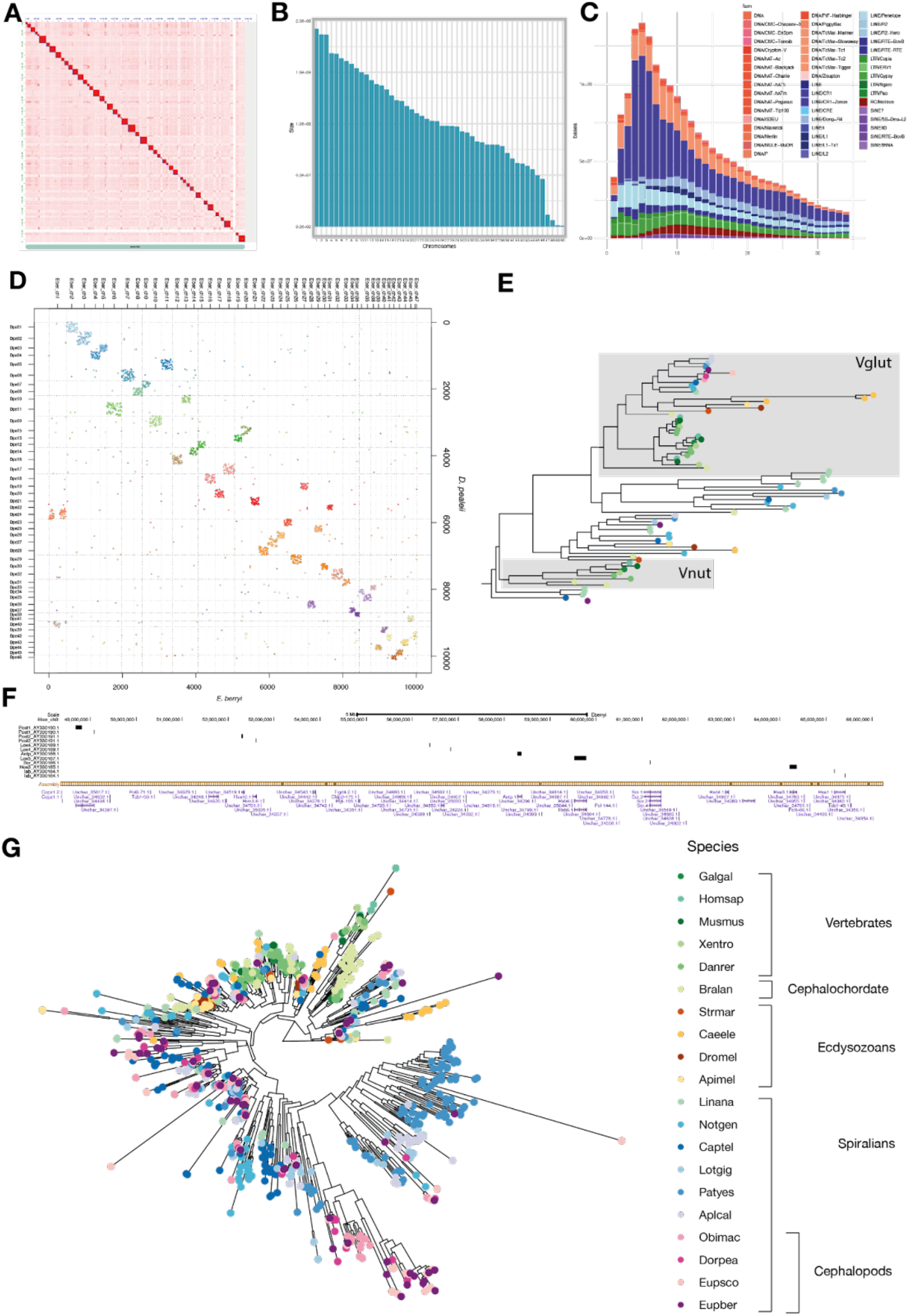
*Euprymna berryi* Genome Assembly QC and Characterisation. (A) Image of HiC contact map (B) Barplot indicating number of chromosomes > 1 Mb sorted by descending length, that were deemed to correspond to a chromosome (C) Repeat landscape (Repeat modeler) (D) Oxford plot comparing chromosome complement of *E. berryi* to that of *D.pealei* (E) Phylogenetic tree of Vglut and Vnut transporters (F) UCSC Genome Browser screenshot demonstrating full 17 Mb Hox cluster in *E. berryi*. (G) Phylogenetic tree of ionotropic acetylcholine subunits across species. Species abbreviations used: Galgal - *Gallus gallus* (chicken), Homsap – *Homo sapiens* (humans), Musmus – *Mus musculus* (mouse), Xentro – *Xenopus tropicalis* (Frog), Danrer – *Danio rerio* (zebrafish), Bralan – *Branchiostoma lanceolatum* (lancelet), Strmar - *Strigamia maritima* (centipede), Caeele – *Caenorhabditis elegens* (nematode), Dromel – *Drosophila melanogaster* (fruit fly), Apimel – *Apis mellifera* (bee), Linana - *Lingula anatina* (brachiopod), Notgen – *Notospermus geniculatus* (nemertean), Captel – *Capitella teleta* (annelid), Lotgig – *Lottia gigantea* (limpet), Patyes – *Patinopecten yessoensis* (scallop), Aplcal – *Aplysia californica* (see hare), Obimac – *Octopus bimaculoides* (octopus), Dorpea -*Doryteuthis peallei* (longfin inshore squid), Eupsco – *Euprymna scolopes* (Hawaiian bobtail squid), Eupber – *Euprymna berryi*

**Figure S2.**
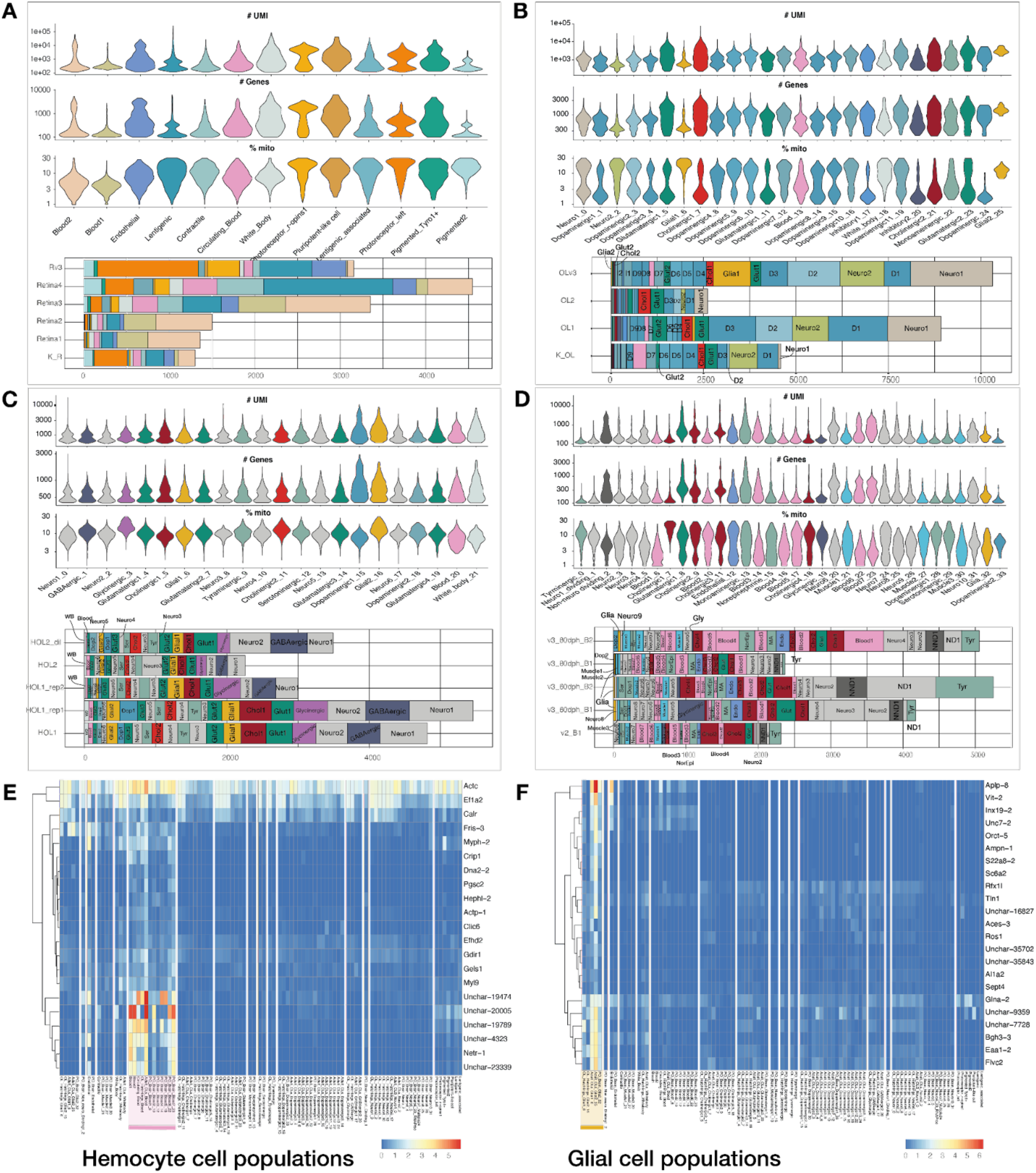
Quality control metrics of scRNA-seq. (A-D) QC Violin plots showing the number of UMIs (top row), genomic features (middle row) and percentage of genes encoded by the mitochondrial genome expressed in each cell cluster (bottom row) and barplots indicating contribution in terms of numbers of cells from each replicate, for each dataset (A) retina (B) adult optic lobe (C) hatchling optic lobe (D) non-optic lobe brain control. (E-F) Heatmap showing non-neuronal markers in scRNA-seq datasets : (E) blood cells, (F) glial cells.

**Figure S3.**
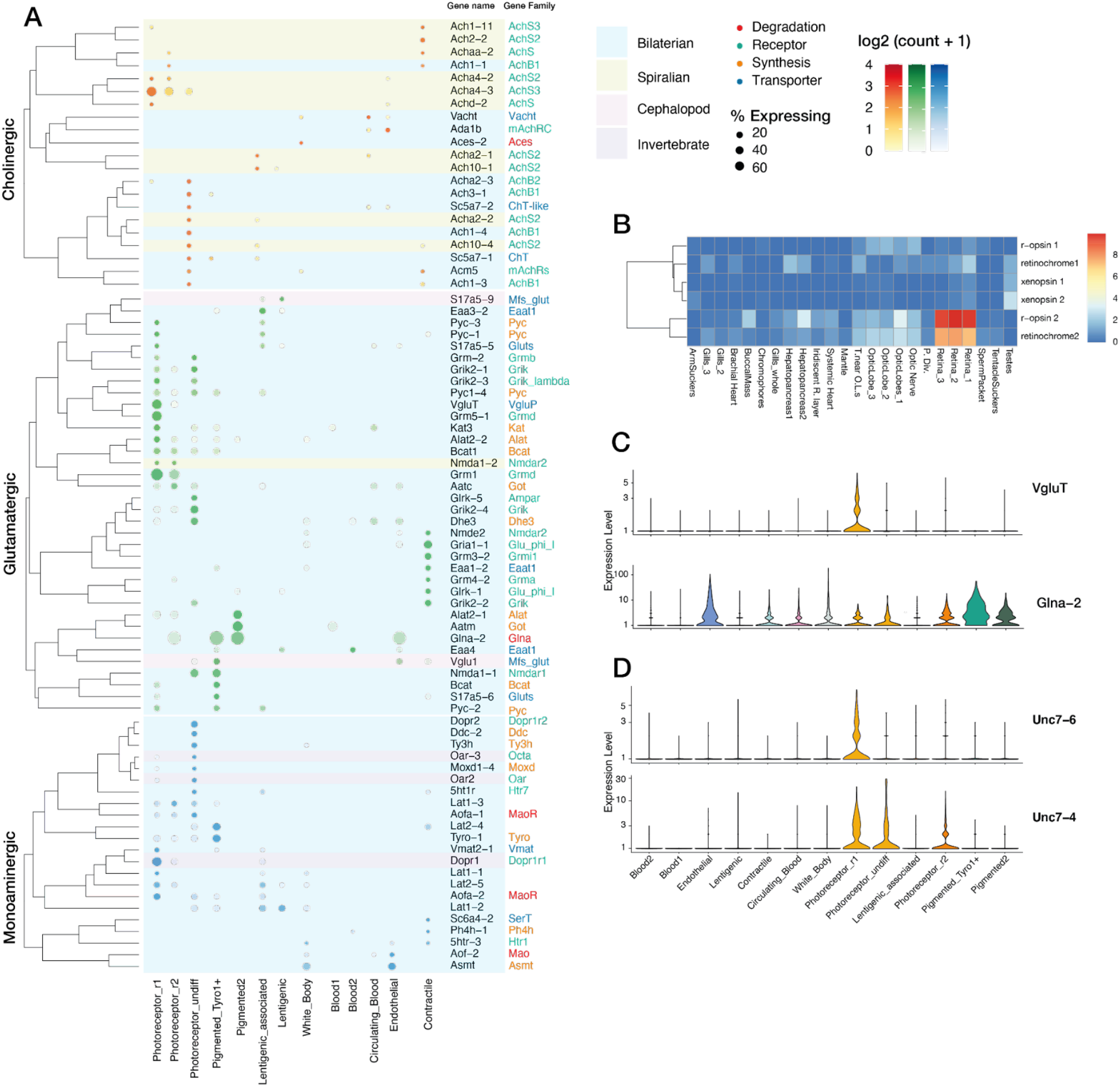
Gene expression in *E.berryi* retina. (A) Dot plot of neuronal markers (in retina scRNA-seq dataset). Circle radius is proportional to fraction of cells in each cluster expressing gene, and color intensity is specified by average scaled expression level as noted. (B) Heatmap of log_10_(FPKM+1) bulk RNA-seq for 6 chromoproteins r-opsin1, r-opsin2, retinochrome-1, retinochrome-2, xenopsin1, xenopsin2 in different organs of a male adult *E. berryi*. (C) Violin plot showing presynaptic glutamatergic marker VgluT and glutamate degrading enzyme Glna-2 expression in retinal scRNA-seq clusters. (D) Violin plots of innexins (gap junction proteins) expression in retinal photoreceptor populations.

**Figure S4.**
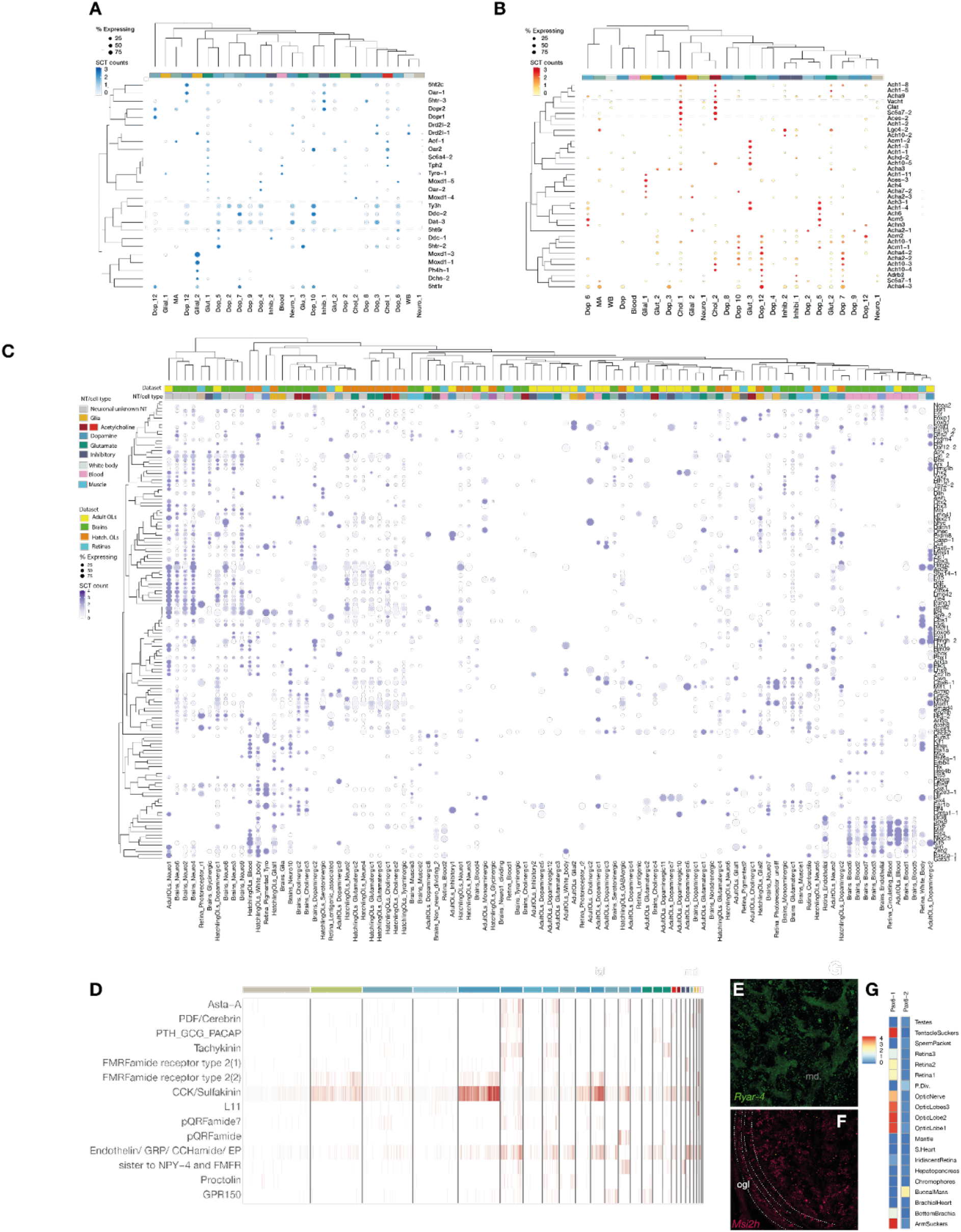
Optic lobe gene expression. (A-B) Dot plots indicating expression levels in the mature adult optic lobes of *E. berryi* of key genes complement: (A) monoamine neurotransmission genes, (B) cholinergic neurotransmitter genes. Box with dotted lines surrounds expression of genes involved in synthesis of neurotransmitters. (C) Dotplot of indicating expression levels of transcription factors in adult and hatchling optic lobe datasets. (D) Heatmap showing expression levels of receptors to neuropeptides in adult optic lobes. (E-G) Fluorescence imaging of HCR for expression of (E) Ryar-4 (RY-amide receptor), (F) Msi2h (Musashi), (G) Heatmap of expression levels of Pax6 paralogs (log_2_(FPKM+1) in bulk RNA-seq on selected organs. Abbreviations used in (A-B): Dop - dopaminergic, Glut - glutamatergic - Chol - cholinergic, WB - white body, MA – monoaminergic

**Figure S5.**
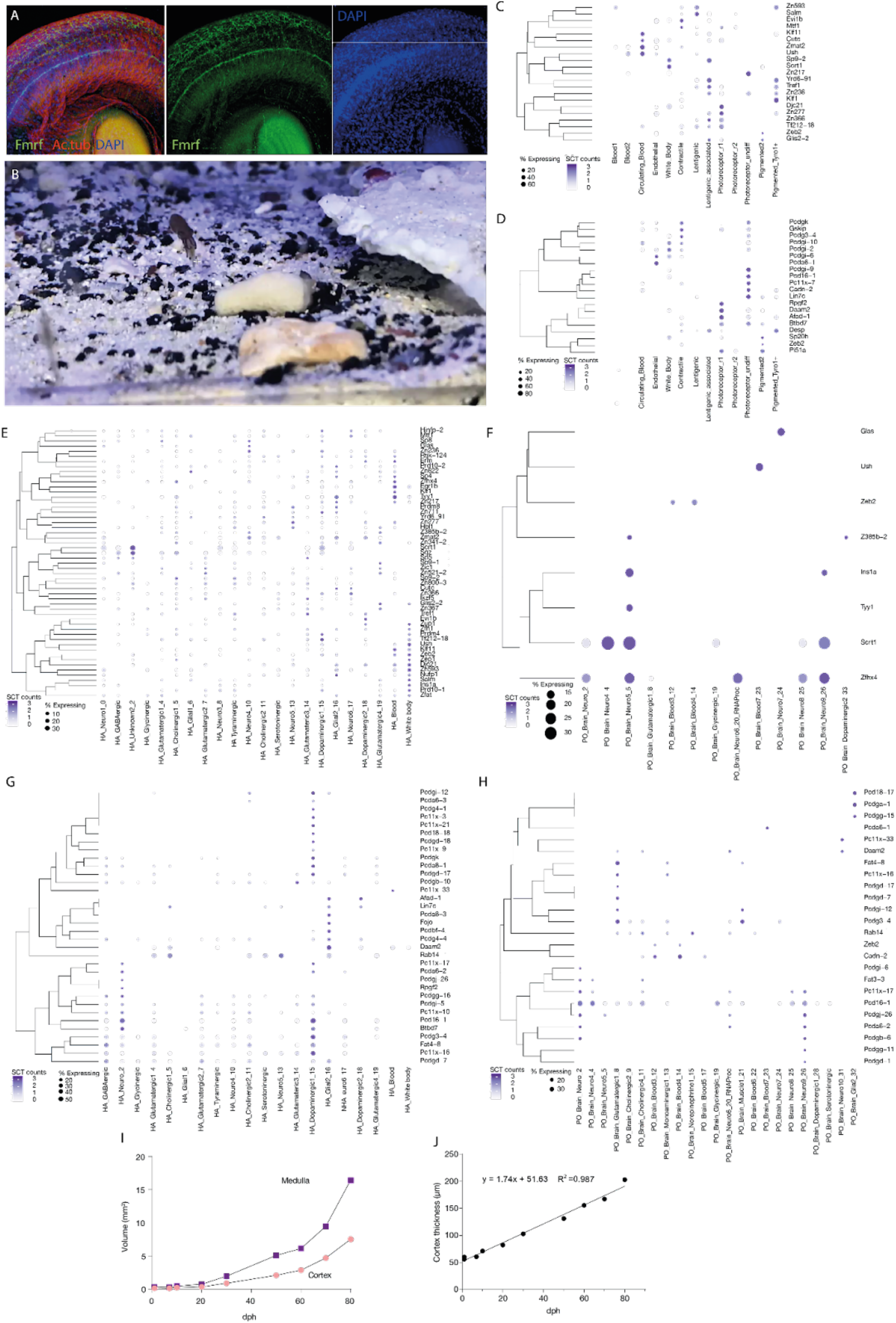
Comparison of gene expression between *E.berryi* retina, hatchling and adult optic lobes. (A) Antibody staining demonstrating the location of FMRF-amide peptide, acetylated tubulin and DAPI in the *E. berryi* retina. (B) Image of *E. berryi* hatchling catching mysid prey (mysid is indicated by cross). (C-H) Dotplots showing expression of protocadherin and C2H2 zinc finger transcription factors, expressed in over 10%(C-F) or over 25% (G-H) of cells of each cell population, in *E. berryi* retina (C-D), hatchling optic lobes (E,G) and perioesophageal brain (F,H): (C) C2H2 zinc finger transcription factor expression in retinas, (D) Protocadherin family expression in retinas, (E) C2H2 zinc finger transcription factor expression in hatchling optic lobes, (F) C2H2 zinc finger transcription factor expression in perioesophageal brain, (G) Protocadherin expression in optic lobes of hatchlings, (H) Protocadherin expression in perioesophageal brain. (I-K) Comparison of optic lobe cortex and medulla over time based on microCT scans. (I) Graph showing the increase in the volume of the medulla and cortex between 1-80dph. (J) Graph showing the increase in the thickness of the optic lobe cortex between 1- 80dph. (K) Rendered 3-D images of segmented optic lobes between 1-80dph with

**Figure S6.**
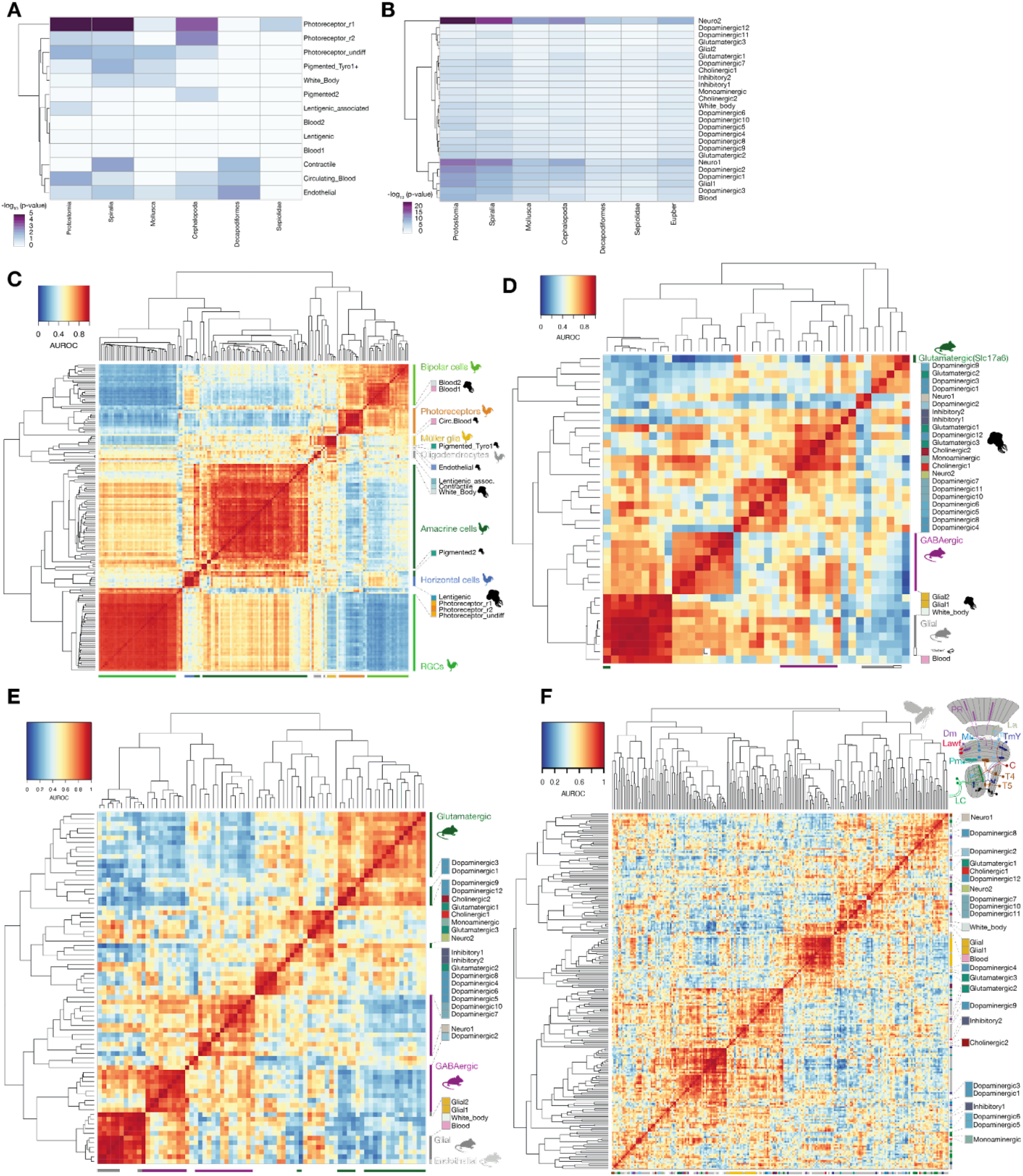
Evolutionary history of genes employed by *E. berryi visual* system. (A,B) Heatmaps showing - log10(p-value) of enrichment test of gene family origin of marker genes for retinal (A) and adult optic lobe (B) cell populations. (C-F) Heatmaps displaying AUROC scores or “Reciprocal_top_hit” match types are identified as cell type pairs comparisons between *E. berryi* retina and chicken retina cell types (Yamagata et al., 2021) (C), *E. berryi* adult optic lobe comparisons to (D-F) to mouse dorsal lateral geniculate nucleus (Allen Brain Atlas, D), to mouse visual cortex (Tasic et al., 2016) (E) and to adult *Drosophila* optic lobes (Özel et al., 2021)(F).

## List of supplementary figures and files

Supplementary Figures:

Figure S1: Euprymna berryi Genome Assembly QC and Characterisation.

Figure S2: Quality control metrics of scRNA-seq

Figure S3: Gene expression in E.berryi retina.

Figure S4: Optic lobe gene expression

Figure S5: Comparison of gene expression between E.berryi retina, hatchling and adult optic lobes.

Figure S6: Evolutionary history of genes employed by *E. berryi* visual system

Supplementary Tables:

Table S1-GenomeAssemblyStats

Table S2-Bulk_RNA-seq_Stages_Organs_mapping_stats

Table S3-HCR_Probes

Table S4-Retina_Cluster_markers

Table S5-AdultOL_Cluster_Markers

Table S6-HatchlingOL_Cluster_Markers

Table S7-Non-OpticLobe_Brain_Cluster_Markers

Table S8-Neuropeptides_in_E.berryi

Table S9-Volumetric_analysis_of_E.berryi_optic_lobes

Table_S10-Evolutionary_origin_and_duplication_of_E.berryi_genes

Supplementary Information:

Video 1: *E.berryi* 2 dph hatchling catching shrimp: IMG_4407.mov

Video 2: *E.berryi* mature adult catching shrimp.

## Materials & Methods

### Data availability

Raw genomic DNA sequencing data and genome assembly generated during this study will be made available through European Nucleotide Archive (ENA) accession number PRJEB52690. Raw and processed bulk-RNA seq and scRNA-seq data will be made available through NCBI’s Gene Expression Omnibus (GEO) accession number GSE203527.

### Ethics statement

This study was carried out in accordance with procedures authorized by Guidelines for Proper Conduct of Animal Experiments by the Science Council of Japan (Science Council of Japan, 2006). Despite the absence of legislation pertaining specifically to cephalopods in Japan, we aspired to abide by the highest standards in the field. All conducted experiments were therefore also in line with EU Directive 2010/63/EU and with the guidelines and the principles detailed in (Andrews et al., 2013; Di Cristina et al., 2015; Fiorito et al., 2015; Smith et al., 2013). All experiments were approved by the Okinawa Institute of Science and Technology Graduate University Animal Care and Use Committee (approval ID: 2018-204). No transgenic animals were used in this study.

### Animal care

Adult *Euprymna berryi* were collected from the coast of Mie prefecture in Japan and transported to Okinawa where they were acclimated to temperature (20°C) and pH (8.3) of a closed aquarium system in filtered natural seawater obtained from the shores of Okinawa, Japan (OIST Seragaki Marine Science Station). Animals were maintained essentially as described previously (Nabhitabhata and Nishiguchi, 2014) until they were sacrificed for experiments. Animals were exposed to a static 12:12 hour light:dark cycle. Tanks that housed the animals contained an enriched environment including natural substrate (autoclaved sand or crushed coral), parts of clay pots and natural rocks as dens. Mature animals were fed daily with Opossum shrimp or mysids, whereas *Neomysis Japonica* proved a suitable prey for hatchlings. Fresh glass shrimp, *Palaemonetes spp*. and frozen shrimp which were purchased in local grocery stores were fed to late juveniles and adults. Tanks were cleaned daily to remove uneaten food and waste matter. Prior to experiments, animals were euthanized using 4% ethanol in sterile-filtered natural seawater. Animals were allowed to breed freely. Hatchlings were obtained either from eggs provided by females impregnated in the wild or by breeding wild animals in the laboratory.

### *E. berryi* genomic DNA extraction and library construction

Germline genomic DNA was obtained from the mature testes of a single mature *E. berryi* male individual by OIST SQC. After thorough grinding in liquid nitrogen, cell lysate was embedded in low-melting agarose plugs, subjected to proteinase K digestion, washed in 20 mM Tris, 50 mM EDTA, pH 8.0 and released using Agarase. DNA molecular size was assessed using a FemtoPulse instrument (Agilent) and found to be made of fragments >50kb suitable for continuous long-read sequencing (CLR). We used several sequencing technologies to assemble the genome. First, two libraries of paired-end 250bp-long Illumina libraries (with insert sizes of 350bp and 800bp) were prepared and sequenced using HiSeq2500 in Rapid Run mode yielding 115Gb and 120Gb Gb providing 19x and 21x coverage respectively at OIST SQC. Demultiplexing was carried out using bcl2fastq (v.2.19). Long-insert Pacific bioscience CLR libraries were prepared using v7 chemistry and 16 SMRT cells were sequenced on a Sequel instrument at the Vincent J. Coates Genomics Sequencing Laboratory at UC Berkeley. Pacbio sequencing yielded 8.8M reads representing 160.7Gb of data with a median read length of 14k and read N50 of 32kb. Finally, we generated 10x linked-reads using the Chromium system (10x Genomics) and sequenced them on 2 lanes of Hiseq4000 in 2×150bp mode (Marks et al., 2019). Demultiplexing was achieved using Supernova (v.2.1.1). Chromatin contact information was obtained from an optic lobe sampled from an adult individual crosslinked in 1% PFA. The HiC library was constructed and the genome was scaffolded using HiRiSE software (Dovetail Genomics)(Koch, 2016) and was sequenced on a HiSeq4000 and NovaSeq6000 SP. Raw sequencing data was submitted to NCBI under accession number PRJEB52690.

### Genome size estimation

The size of the genome was estimated using a k-mer spectrum approach as described in(Ferguson et al., 2014) from 17-mer to 31-mer distribution calculated by Jellyfish (v.2.2.7) (Marçais and Kingsford, 2011). By performing a sum of unique k-mers weighted by their multiplicity, we originally found an effective haploid genome size of 5.6Gb.

### Genome assembly and validation

We used wtdbg2 (v.2.5) with default parameters and genome size parameter “-g 4.6G” to assemble the Pacific bioscience reads (Ruan and Li, 2020). To generate consensus sequence contigs, Pacific bioscience reads were mapped with minimap2 (v.2.16)(Li, 2018) and to the obtained contig assembly in two rounds using the wtdbg2 polishing module. We also utilized the Illumina paired-end date in an additional polishing step using Racon (v.1.3.2) (Vaser et al., 2017), resulting in a haploid reference of 5.9Gb composed of 51,130 contigs and an N50 of 831,242 bp. We further scaffolded contigs with scaff10x (https://github.com/wtsi-hpag/Scaff10X) with parameters ‘ -longread 1 -gap 100 -matrix 2000 -read-s1 10 -read-s2 10 -score 20 -edge 50000 -link-s1 10 -link-s2 10 -block 50000‘. We assessed the completeness and haplotype collapse of the assembled genome using BUSCO (v3.1.0) (Simão et al., 2015), yielding C:84.9% [S:82.7%,D:2.2%], F:5.8%,M:9.3% n:978. (Fig.S1) We also verified the homogeneity of base coverage in final contigs by remapping both the Pacbio using minimap2 (Li, 2018) and Illumina datasets using BWA (Li and Durbin, 2009) and analyzing the BAM alignments with the alfred tool (Rausch et al., 2019) (https://github.com/tobiasrausch/alfred). BAM files were sorted using Samtools (v.1.9)(Li et al., 2009).

To further extend the contiguity of our haploid reference, we used long-range contact information from HiC. The assembly was scaffolded using the HiRise pipeline by Dovetail Genomics. To validate the output assembly, HiC data was processed using Juicer (v.e0d1bb7) and contact density was examined and inspected using Juicebox software (Durand et al., 2016). Our final assembly shows 48 main chromosomes (length > 1Mb, Figure S1A-B). The number of chromosomes is consistent with the reported diploid number of chromosomes 96 in decapods (Jazayeri et al., 2011; Wang and Zheng, 2017). Additional quality control consisted of re-mapping Illumina RNA-seq data and Pacific biosciences Isoseq reads (see below). Bulk RNA-seq Illumina reads contained on average 86.58% of reads mapping uniquely to the genome. Mapping Iso-seq reads yielded 50’400 aligned (99.51%), of which 45,243 on 100% of their length, 47,379 at 95% and 48,829 at 75% (i.e. 97% of transcripts).

### Bulk RNA extraction and sequencing

We obtained total RNA from embryos over embryonic developmental (Stage 0 - 28) and from 26 samples of organs of a >100-day old adult male. Total RNA extraction was carried out using TRIzol reagent (Invitrogen) for lysis followed by on-column purification using PureLink RNA Mini Kit (Invitrogen) as described in TRIzol Plus RNA Purification instructions from manufacturer (https://assets.fishersci.com/TFS-Assets/LSG/manuals/Trizol_Plus_man.pdf). Briefly, tissues were dissected and directly homogenized in 1 mL of TRIzol Reagent on ice. After 5 minute incubation at room temperature, 0.2mL of chloroform was added, vortexed and incubated for an additional 3 minutes at room temperature. Phase separation was achieved by centrifugation at 12,000g at 4°C. The upper colorless phase was mixed with an equivalent volume of 70% ethanol in DEPC-treated water, mixed, vortexed and applied directly to the PureLink RNA Mini Kit columns. Wash steps were followed as per manufacturer’s protocol. RNA extraction was followed by Turbo DNase treatment (Thermo Fisher Scientific) to remove genomic DNA according to manufacturer’s instructions. RNA quality was evaluated using Tapestation 4200 BR RNA Screentape and reagents (Agilent Technologies). For organ-specific transcriptomes, library preparation was achieved by using TruSeq Stranded mRNA Library Prep for NeoPrepTM (Illumina). For stage-specific transcriptomes, we used NEBNExt Ultra II Directional RNA Library Prep Kit for Illumina (NEB). Organ-specific RNA-seq was carried out on a Hiseq4000 and embryonic stage specific RNA-seq was done on a Novaseq6000 SP, yielding a total of 1,966,727,071 reads. Demultiplexing was carried out using bcl2fastq (v2.19). Reads were mapped to the *E. berryi* genome using STAR v2.5.2 in several rounds (Dobin et al., 2013). In a a first round, the following parameters were used: ‘--outSAMmapqUnique 255 --outFilterMultimapNmax 10 --outReadsUnmapped Fastx --outFilterMismatchNmax 999 --winBinNbits 10 --chimOutType SeparateSAMold -- chimSegmentMin 20 --chimJunctionOverhangMin 20 --outFilterMismatchNoverLmax 0.5 -- outFilterMismatchNoverReadLmax 10 --outSAMstrandField intronMotif -- alignSoftClipAtReferenceEnds No --alignMatesGapMax 5000 --outFilterScoreMin 100’. In a second round, we utilized the junctions observed in the first round, using the following parameters: --outSAMstrandField intronMotif --outSAMtype BAM SortedByCoordinate -- outSAMmapqUnique 255 --outFilterMultimapNmax 1 --outReadsUnmapped Fastx -- chimJunctionOverhangMin 20 --alignSoftClipAtReferenceEnds No –sjdbFileChrStartEnd *SJ.out.tab --limitSjdbInsertNsj=5000000 --sjdbInsertSave All’. Count tables were obtained using subreads FeatureCounts(Liao et al., 2014) using ‘-p -B -t exon -g gene_id ‘ parameters. We also sequenced full-length cDNA from retinas and optic lobes on a Sequel instrument. Following the Isoseq protocol, circular consensus (CCS) of subreads were calculated, and subsequently classified as full-length or non full-length based on the presence of SMART adaptors at both extremities. Full-length transcripts were clustered and polished using all ccs reads with quiver.(https://github.com/ben-lerch/IsoSeq-3.0).

### Genome annotation

We generated RNA-seq data for adult organs and developmental time series with 1,966,727,071 reads. (Table S2-3). We aligned the reads to the genome using STAR (v2.5.2b) and reached > 88 % uniquely mapping reads (Dobin et al., 2013). These alignments were subsequently used to assemble transcriptomes for each organ using Stringtie (v1.3.3b) (Kovaka et al., 2019; Pertea et al., 2015, 2016) and then merged together using Taco (Niknafs et al., 2017). In parallel, a *de novo* assembly of all RNA-seq was performed using Trinity (Haas et al., 2013). Assembled transcripts from *de novo* and genome-guided Trinity, as well as the high-quality Isoseq transcripts were aligned to the genome using GMAP (version of 2019-02-26)(Wu and Watanabe, 2005) using parameters ‘-f 3 -n 0 -x 50 -t 16 -B 5 --gff3-add-separators=0 --intronlength=500000’. Mikado (v1.2.1)(Venturini et al., 2018) was used to generate a high-quality reference transcriptome leveraging (i) the trinity transcriptomes, (ii) the Iso-seq transcripts, (iii) the Stringtie transcriptomes merged with Taco and a set of curated splice-junctions generated from RNA-seq alignments using Portcullis (v1.0.2) using parameters ‘full --strandedness firststrand --bam_filter’ (Mapleson et al., 2018). Putative fusion transcripts were detected by Blast comparison against Swissprot and ORFs were annotated using Trans-decoder (https://github.com/TransDecoder/TransDecoder). Transcripts derived from the reference transcriptome were selected to train Augustus (v.3.3.3) *de novo* gene prediction tool (Stanke et al., 2006a). Exon positions in the Mikado transcriptome assembly were converted into hints for Augustus gene prediction. We aligned the proteomes of cephalopod species bobtail *Euprymna scolopes (Belcaid et al., 2019)*, octopod *Octopus bimaculoides (Albertin et al., 2015b)*, squid *Doryteuthis peallei (Albertin et al., 2022c)* and provided them as CDS hints for Augustus gene prediction.

Finally, a repeat library was constructed using RepeatModeler (v.1.0.11)(Smit and Hubley, 2008) and used for masking with RepeatMasker (v.4.0.7)(Nishimura, 2000). Gene models with more than half or more of their exons overlapping with more than half of those of repeats were discarded, yielding 56,767 filtered gene models. Alternative transcripts and UTRs were subsequently incorporated using the PASA (v.2.41) pipeline (Haas et al., 2008). PFAM domains were identified using PfamScan (v.1.6) The obtained gene models contain a total number of 5,277 distinct PFAM domains (SuppData-PFAM domains).

### Gene family reconstruction and phylogenetic analyses

We reconstructed gene families using Orthofinder (v2.33) using the non-redundant protomes of selected species ranging across the tree of bilaterian and including some key model species for neurobiology work such as fruit fly, mouse and zebrafish (Emms and Kelly, 2019). We recovered 21,332 gene families, of which 12,046 contained at least one *E. berryi* gene family member.

### Dissociation of tissues into single cells

Tissues were dissociated using a 1% Papain solution in filtered natural seawater containing 0.1% sodium thioglycolate for retinas, and a 1% Pronase solution for non-retinal tissues on a nutator with gentle trituration. Cell clumps were filtered out using either 30µm MACS SmartStrainers (Miltenyi Biotec) for v2 sample or using 40µm and 70µm cell strainers (flowmi). Cells were resuspended in a 50% L15 medium in sterile-filtered natural seawater. Single cell suspension quality and cell concentration was evaluated using C-Chip Disposable hemocytometer (NanoEntek) on a Zeiss PrimoVert Monitor Inverted microscope. Number of viable cells was evaluated using Trypan blue (Thermo Fisher).

### Single cell library preparation and sequencing

scRNA-seq library preparation was performed immediately after dissociation with the 10X Chromium v2 or v3.1 kits (10X Genomics) following manufacturer’s protocol. We produced 5 replicates of optic lobes scRNA-seq libraries from *E. berryi* that had hatched the same day (1dph), and 4 replicates of optic lobe scRNA-seq libraries from mature adult *E. berryi* (1x>80dph, 1×60dph, 2×80dph). We were unable to produce libraries derived from retinas of 1dph hatchlings and used 6 replicates of adult individuals (2×60dph, 4×80dph, 1×> 80dph). We dissected the perioesophageal brain without the optic lobes to use as a control from mature adult *E. berryi* and generated 5 independent libraries (2x 60dph, 2×80dph, 1x >80dph). (STable4, Fig.S2). scRNA-seq was carried out on either an Illumina HiSeq2000, HiSeq4000 and NovaSeq6000 instrument at OIST SQC. The samples obtained from > 80dph-old individual produced using the v2 kit, were originally sequenced on a Hiseq4000 paired-end 150bp run with scattered PhiX spikes ranging from 0 to 40% on each lane to reduce base bias. Raw sequencing data was processed with the cellranger version 2.10 and subsequently v.3.1.0 mkfastq script (10x Genomics). Re-sequencing of the same libraries was achieved using an Illumina Novaseq6000 instrument using paired-end 150bp with 10% PhiX spike, and was similarly demultiplexed using cellranger mkfastq v.3.1.0. The paired end 150bp-long fastq files were trimmed by quality and for size using sickle (https://github.com/najoshi/sickle). Read quality was examined using fastqc (v.0.11.5). All the other single cell libraries were sequenced using the following program: Read1:26bp, Index1:8bp, and Read2:98bp and demultiplexing was achieved using bcl2fastq (v.2.19). Hatchling (1dph) optic lobe samples were sequenced on a Novaseq6000 instrument (Illumina) using a 10% PhiX spike.

### scRNA-seq read alignment and analysis

For scRNA-seq mapping only, all gene models were extended by 3kb in the 3’ direction using the script https://github.com/JulienPPichon/UTR_extension_GTF. Reads were mapped to the genome with STAR Solo (Bolger et al., 2014; Dobin et al., 2013). (STable4, Fig.S2). A total of 6,709,213,691 reads were sequenced, of which 5,510,308,930 mapped uniquely (corresponding to 82% of all sequenced reads) (Table S4). STAR Solo based gene-barcode counts matrices were analyzed with the Seurat (v.4.1.0) R package (Hao et al., 2021). Numbers of features, UMIs and percentage of reads mapping to the mitochondrial genome per cell were examined for each dataset (Figure S2A-D). Features found in less than 3 cells were removed from subsequent analysis. Cells in which more than 30% of reads corresponded to genes encoded by the mitochondrial genome were removed from subsequent analysis. A minimal number of cells expressing each feature was set for each dataset (TableS4) ranging from 100 to 600 features per cell. Replicates were normalised individually using ‘SCTransform()’ function prior to integration. Using ‘PrepSCTIntegration()’ and ‘FindIntegrationAnchors()’ with ‘normalization.method’ set to ‘SCT’. Data integration was achieved using the ‘IntegrateData’ command. Principal component analysis was performed using the ‘RunPCA()’ command using 1:200 principal components at first. The ‘FindNeighbors()’ and ‘FindClusters()’ commands were used to identify putative clusters at resolutions of 0.1, 0.2, 0.5, 0.75, 1, 2, 5 and 7. The number of relevant principal components was determined using the ‘ElbowPlot’ function of Seurat and by examining gene expression in UMAPs created using principal components from 1:10-200. We used the Seurat’s ‘FindAllMarkers’ command and R package presto’s ‘wilcoxauc’ command to identify differentially expressed genes in each cluster. In a first time, using only the >80dph and 60dph samples of the retinal datasets, using 200 principal components with a resolution of 0.1, a group of cells that clustered with all other clusters across resolutions was observed (‘cluster6’ at resolution=0.1 corresponding to 74 cells) and was removed as this behavior was reminiscent of cell doublets. In a second time, after the other retinal datasets were generated and re-clustered using 1:35 principal components and an additional group of cells that expressed markers of both neurons and hemocytes (cluster 2 at resolution=0.2, corresponding to 2,488 cells) was deemed to correspond to cell doublets and removed from subsequent analysis. For the retina dataset, we utilized higher resolution cell clusters for neuronal and retinal clusters using resolution 0.5 for the photoreceptors, pigmented-tyro+, pigmented2, Pluripotent-like cells, white body and lentigenic-associated cell populations, and resolution 0.2 for lentigenic cells, but a lower resolution of 0.1 for clusters representing cell types that are not specific to the retina (hemocytes, endothelium, contractile cells), as our interest does not lie in these cell types. For the adult optic lobe dataset, we used 200 principal components and a resolution of 0.75. For the hatchling optic lobe dataset, we used 200 principal components and a resolution of 1. For non-optic lobe brain control datasets, we used a resolution of 3. Markers of each cluster for each dataset are available in Tables S6-9.

### X-ray microtomography (micro-CT)

Euthanized animals were fixed in 4% PFA in seawater for at least overnight at 4 °C. Samples were dehydrated serially in 25%, 50%, 75% and 100% ethanol at 4 °C. Samples were kept in each concentration of ethanol overnight and then transferred to 1% iodine solution (1.9g iodine per 50ml of absolute ethanol) at room temperature for at least 24 hours (or longer for larger samples). After staining, samples were washed in absolute ethanol three times.

X-ray microscope Xradia 510 Carl (Zeiss) was used to obtain microCT images of squid samples. Small hatchlings were mounted in heat-sealed pipette tips or 0.2 ml PCR tubes. Older samples were mounted in 1.5ml Eppendorf tubes. In the containers used during image acquisition, samples were either submerged in ethanol or placed on Kim wipe or plastic wrap soaked in ethanol. Kim wipe and saran wrap was also used to prevent samples from moving within the container. We used standard objectives (0.39x, 4x) and different combinations of source-sample and sample-detector distances to fit and optimize the entire region of interest (S**Table 1**). Images were acquired with an exposure time of 1-2 seconds per projection and a total of 1601 projections over 360 degrees. No filter was used. The vertical stitching feature was applied to join multiple tomographies.

The projection data from each scan was reconstructed using the integrated volume reconstruction software of the Xradia machine. The resulting reconstructions were exported in .txm or DICOM format and imported into Amira (version 6.5, Thermo Fisher Scientific, Waltham, Massachutsetts, USA) for manual segmentation of anatomical structures and to render isosurfaces. For CNS brain lobes, we segmented the inner, lobular neuropil without its confluent outer perikaryal layer. The neuropil layer in the center of lobes can be seen as darker shades of gray in tomography, while the surrounding perikaryal layer is seen as lighter shades of gray (Fig 1F).

The surface generation algorithm implemented in Amira (*Generate Surface* module) was applied for each segmented material using smoothing function values < 2.5. The resulting 3D surface meshes were exported in .ply format and imported into ParaView package for data visualization (Ayachit, U. (2015). *The paraview guide: a parallel visualization application*. Kitware, Inc..). All figures were created with Adobe Illustrator (Adobe Inc., 2019., available at: {https://adobe.com/products/illustrator)}.

#### Transmission electron microscopy

Optic lobe and retinal samples dissected from freshly sacrificed animals were fixed in 4 % PFA in filtered seawater overnight at 4°C. Samples were then stored in 2.5 % glutaraldehyde in 1x PBS for up to 1 month at 4 °C. To enhance contrast of the samples for TEM observation, samples were stained with 2% osmium tetroxide in water for 1 hour and then washed with deionized water 5 times for 3 minutes each. Samples were then blocked stained in 4% uranyl acetate overnight at 4 °C, and washed with deionized water 4 times for 3 minutes each. Samples were then serially dehydrated in 30% and 70% acetone diluted in water for 15 minute each. Samples were then further dehydrated in 100% acetone for 15 minutes.

The 15-minute dehydration step in fresh 100% acetone was repeated two additional times. For embedding samples, epoxy consisting of 20ml Epon 812 Resin, 10ml DDSA EM, 10ml MNA (hardener) and 1ml DMP30 were mixed in a 50ml conical tube and then rotated in a rotor for 1 hour at before used. Samples were first incubated in 33% and then 77% epoxy diluted in 100% acetone for 1 hour each, and then transferred into 100% epon for 1 hour-incubation. Samples were then transferred into fresh 100% epon for overnight incubation. All steps that are part of the acetone dehydration and epon incubation up to this stage were performed at room temperature.

To embed samples in epoxy, samples were placed in a silicon mould filled with epoxy resin Epon formula (Taab Company, UK) and then incubated at 60 °C for 2 days in an oven. After polymerization of the epoxy, blocks with samples were sliced into 70 nm sections using a Leica EM UC7 Ultramicrotome and placed on grids (Nisshin). For sectioning, a diamond knife (Diatome Ltd, Switzerland) was used. Before observation under a transmission microscope, grids with sections were re-stained with 4% uranyl acetate for 20 minutes and then washed with deionized water three times for 3 minute each. Image acquisition was performed with a Transmission Electron Microscope (JEOL JEM-1230R, 100 KV TEM). Image acquisition parameters are listed in STable1.

### Hybridisation chain reaction (HCR) stainings

*E. berryi* were washed several times in natural seawater, and euthanised in 4% ethanol in sterile-filtered natural seawater, and fixed in 4% formaldehyde in sterile-filtered natural seawater overnight at 4°C. After 3 washes in PBSx1, squid were dehydrated in increasing concentrations (25%:75%,50%:50%, 75%_25%,100%,0%) of methanol:PBS for 15 minutes each and stored in 100% methanol at −30°C until necessary. For staining, individuals were rehydrated in a reversed-order methanol-to-PBS series (25%:75%,50%:50%,75%_25%). For wholemount stainings, rehydrated individuals were utilized directly for HCR. When cryosections were required, fixed samples were first incubated in 25% sucrose in PBS solution overnight, and subsequently in a 35% sucrose in PBS solution overnight or until the sample sank to the bottom of the falcon tube. After sucrose incubation, samples were embedded in OCT cryosection medium and cryosectioning was performed using either a CM3050S or CM1950 cryostat (Leica). HCR probes (version3 chemistry) were designed using the e Molecular Instruments website (www.molecularinstruments.com). RNA sequences utilized for probe design are available in Table S5. Section HCR staining was briefly described as below: slides were rinsed in chilled 1XPBS for 1h at 4°C to remove OCT, then enclose the samples with iSpacer (0.5mm, SUNJin Lab), surround inside the iSpacer square with pap pen. Slides were transferred to a humidified chamber at 37°C and treated with 10ug/mL Proteinase K in PBST for 10 min to permeabilize cell membrane, followed by two rinses with 2mg/mL glycine in PBST at room temperature(RT) and post-fixation with 4% PFA for 30min. Sections were washed with PBST for 5 min 3 times before pre-hybridization. Each sample was pre-hybridized by adding 200μl of hybridization buffer and incubated at 37°C for 10 min inside the humidified chamber. Hybridization was carried out by exchanging the hybridization buffer with 0.4pmol of probe set, followed by overnight incubation at 37°C. After overnight incubation, excess probes were sequentially rinsed in probe wash buffer to which 5XSSCT had been added to final concentrations (vol/vol) of 0%, 25%, 50%, 75%, and then 100% for 15 min each at 37°C. Slides were rinsed before pre-amplification with 5XSSCT for 5 min, and incubated in amplification buffer for 30 min at RT. After removing the pre-amplification buffer, 100μl of hairpin solution, prepared according to the manufacturer’s instructions and snap-cooled, was added to each sample and incubated in a dark humidified chamber at RT overnight. Excess hairpin solution was removed through two rinses with 5X SSCT for 30 min, followed by a final rinse for 5 min. Finally, ProLong Gold Antifade Mountant (ThermoFisher) was added to the samples, and sections were sealed with coverslip for imaging. Whole mount HCR was carried out by following the manufacturer’s protocol (whole-mount sea urchin embryos’ protocol), with an additional clearing step using the RapiClear solution (SUNJin Lab). Samples were mounted in an iSpacer (1mm, SUNJin Lab) filled with RapiClear solution. Staining results were visualized using LSM780/LSM880 at OIST Imaging facility or LSM980 at UCL Imaging Facility. Images were analyzed using ZEN software and Fiji.

### Comparative evolutionary analysis

Datasets count tables and embeddings were downloaded from https://singlecell.broadinstitute.org/single_cell for the following datasets. Seurat objects were generated according to the embedded cell annotation. Datasets were reduced to genes belonging to shared gene families for both species compared and merged (Crow et al., 2018; Wang et al., 2021). After determining variable genes, Aurocs scores were calculated using the Unsupervised metaneightbour function in fast mode and normalizing votes.

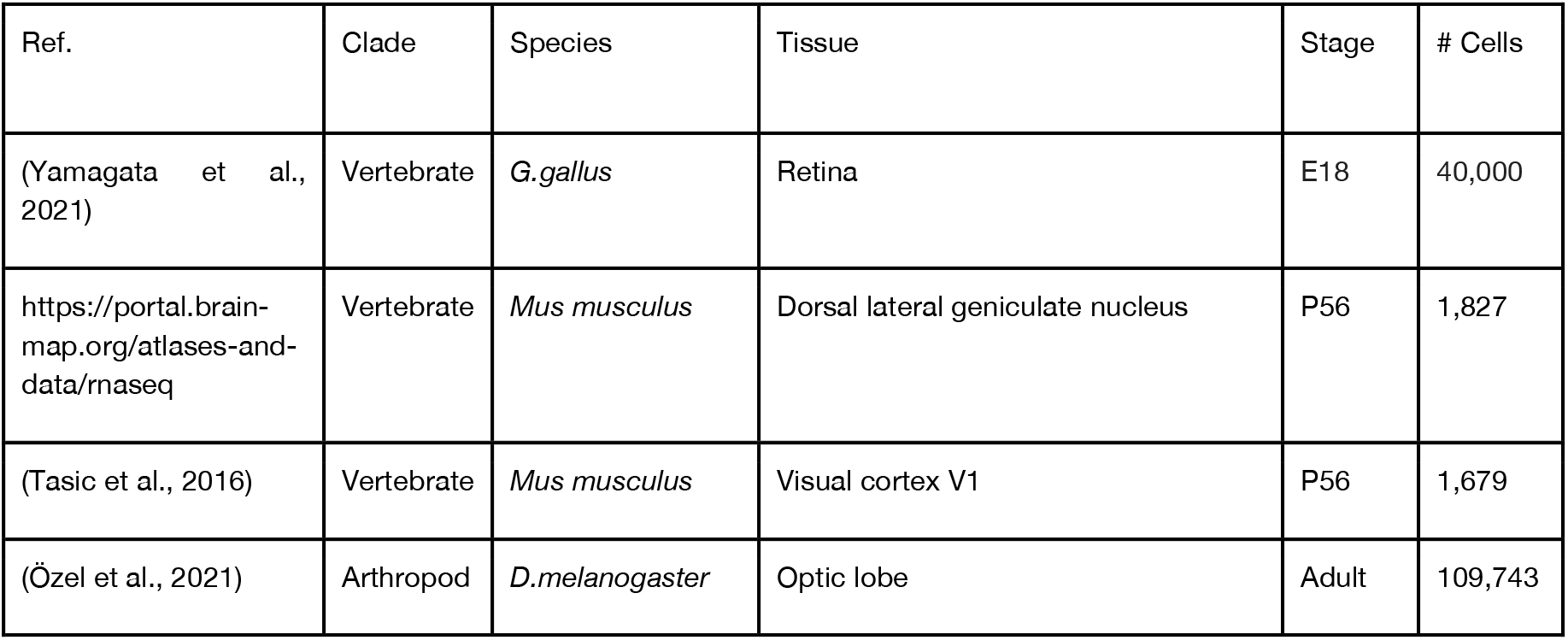

## Acknowledgements

We acknowledge the OIST sequencing facility ‘SQC’ and its members for their support: N. Arakaki, S. Yamasaki, M. Kawamitsu, T. B. H. Soliman, as well as Vincent J. Coates Genomics Sequencing Laboratory at UC Berkeley and particularly S. McDevitt for Pacific Bioscience sequencing. OIST High-throughput computing facility servers were used for computational analysis. We owe special thanks to T. Kakeya from Scrum Inc. for teaching us to perform scRNA-seq libraries using 10x Genomics reagents. We thank I. Masai for access to cryostat and Y. Nishiwaki for assistance and training to utilize the cryostat, and M. Hall and A. Takahashi from the OIST Imaging section for assistance with cryosectioning. We thank A. Greig from the UCL Imaging facility for expert advice on imaging and training and A. Sadier for advice on HCR stainings, and are grateful to T. Sasaki and M. Hall from OIST Imaging Section for training and assistance with TEM. We thank the Evolutionary Neurobiology (Watanabe) Unit at OIST for the anti-Fmrf-amide antibody. We thank R.Nakajima for professional photography of Euprymna berryi, and J. Jolly and R. Kawaura for early contributions to bobtail squid aquaculture. We thank J.Pichon for optimizing the UTR extension python script. We are grateful to C. Plessy, C. Martín-Durán, A. de Mendoza, M. Feller, K. Shekhar and M. Kuba for critical reading of the manuscript and helpful comments and suggestions. We are grateful to Z. Ziadi for insightful calculations based on microCT and optic lobe growth modeling. This work was supported by internal funds of the OIST Molecular Genetics Unit, the Chan Zuckerberg Biohub to D.S.R. D.G. was supported by a JSPS fellowship for prospective researchers (Fellow ID No. PE17011). D.S.R. is grateful for the support of the Marthella Foskett Brown Chair in Biology at UC Berkeley. F.M. was supported by a JSPS Kakenhi grant (#19K06620), BBSRC and Royal Society URF funding.

## Author contributions

D.G., F.M. and D.S.R. conceived the project. D.G. and F.M. assembled the genome, carried out bulk RNA-seq extractions, performed annotation, performed scRNA-seq and analyzed data. Y.T. and Y.H. carried out histology and HCR and antibody stainings. F.K.-Z. And Y.H. carried out micro-CT sample preparation. C.S., Y.H. and F.K.-Z. analyzed micro-CT. L.Z. took care of animals, extracted hemocytes, prepared samples for histology and prepared cryosections. C.S. oversaw animal culture and helped with dissections. C.S. and Y.T. carried out photography. L.P. carried out imaging of HCR stainings. N.L. contributed to staining, imaging effort and supervised work. D.G., F.M. and D.S.R. wrote the manuscript with input from all authors.

